# Structures of the stator complex that drives rotation of the bacterial flagellum

**DOI:** 10.1101/2020.05.12.089201

**Authors:** Justin C. Deme, Steven Johnson, Owen Vickery, Amy Muellbauer, Holly Monkhouse, Thomas Griffiths, Rory Hennell James, Ben C. Berks, James W. Coulton, Phillip J. Stansfeld, Susan M. Lea

## Abstract

The bacterial flagellum is the proto-typical protein nanomachine and comprises a rotating helical propeller attached to a membrane-embedded motor complex^1^. The motor consists of a central rotor surround by stator units that couple ion flow across the cytoplasmic membrane to torque generation. Here we present the structures of stator complexes from multiple bacterial species, allowing interpretation of the extensive body of data on stator mechanism. The structures reveal an unexpected asymmetric A_5_B_2_ subunit assembly in which the five A subunits enclose the two B subunits. Comparison to novel structures of other ion-driven motors indicates that this A_5_B_2_ architecture is fundamental to bacterial systems that couple energy from ion-flow to generate mechanical work at a distance, and suggests that such events involve rotation in the motor structures.

## Main

A motor is a machine that supplies motive power for a device with moving parts. Biological systems use both linear and rotary motors to generate a variety of outputs. One of the most fascinating and complex biological rotary motors is the flagellar apparatus used by bacteria to propel themselves through fluid environments. Although bacterial swimming was first observed in the 17^th^ century^2^, a mechanistic understanding of how the bacterial flagellum generates rotation is still lacking. The core of the flagellum is a highly conserved motor (Fig. 1a) consisting of a cytoplasmic-membrane embedded rotor complex surrounded by varying numbers of stator complexes (hereafter termed simply stators) that generate torque^3^. While high resolution information has recently been obtained for the rotor component^4^, structural detail of the stators has thus far been limited to modelling studies^5^.

**Figure 1.**
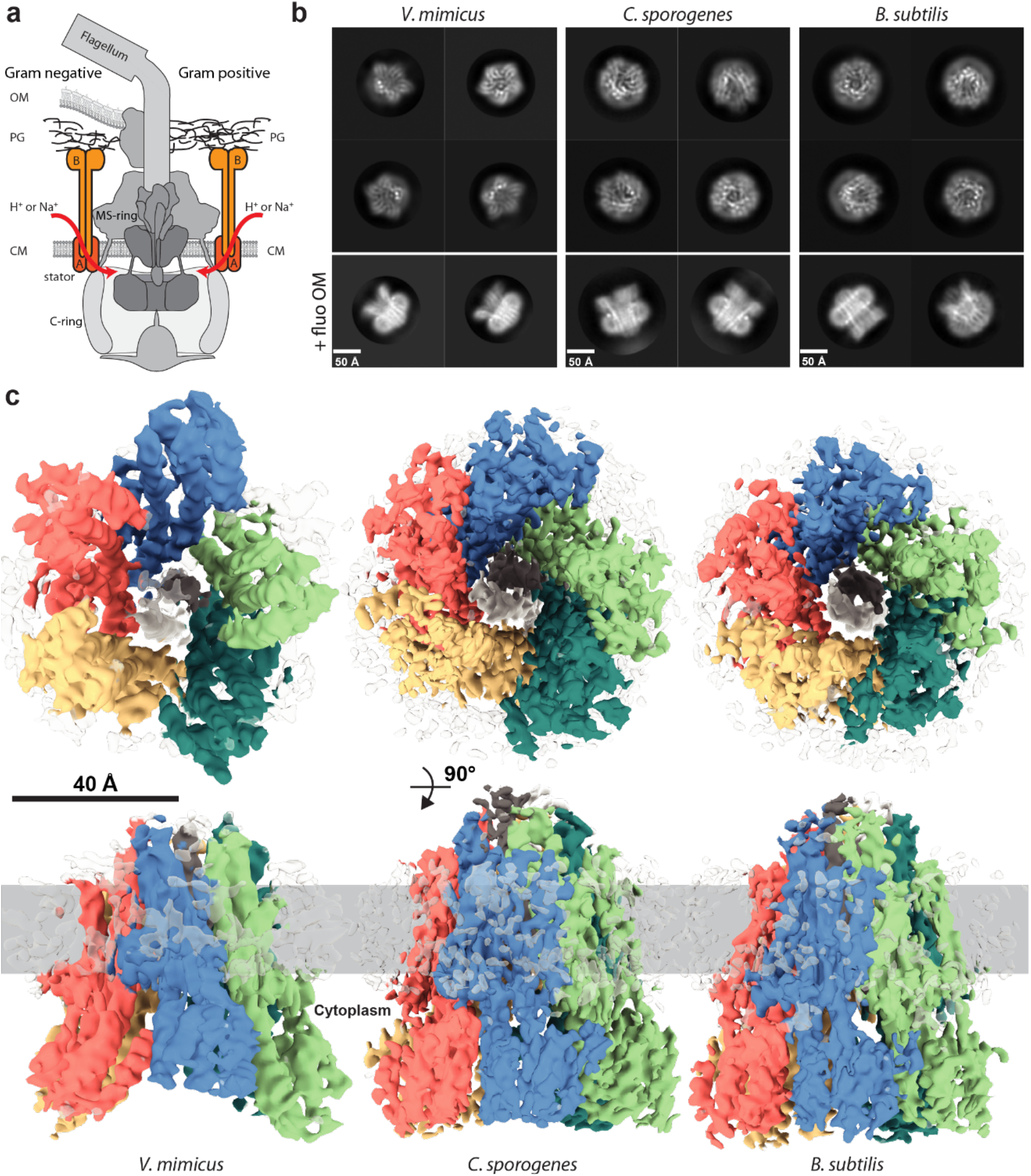
Stators from multiple organisms have a MotA_5_MotB_2_ stoichiometry. **a**, Composite cartoon showing the general organisation of bacterial flagellar complexes in Gram-negative (left side) and Gram-positive (right side) bacteria with major components labelled. Stators are orange and rotor components, MS- and C-ring, are grey. OM, outer membrane; CM, cytoplasmic membrane; PG, peptidoglycan. **b**, 2D class averages of cryo-EM particles of stators from the bacterial species indicated. Upper panels are representative ‘top’ views of the 5:2 complexes. Lower panels are ‘side’ views from data collected in the presence of fluorinated octyl maltoside. **c**, Cryo-EM volumes of stators from the three bacterial species. The MotA subunits are coloured pink, blue, green, teal, and yellow, and the centrally-located MotB subunits white and dark grey. Bound detergent is shown as transparent density at the periphery. Upper panels show views from the cytoplasm, lower panels show side views with the likely membrane location (assigned from the position of the detergent micelle and from simulations; Extended Data Fig. 6) indicated by the grey bar.

Stators harvest energy from either H^+^ or Na^+^ ion flow across the cytoplasmic membrane, generating torque in the cytoplasmic portion (C-ring) of the rotor complex^6–9^. Chimeras between H^+^- and Na^+^-dependent stators are functional, implying that the mechanism converting ion flow into work is the same for the two coupling ions^10^. Stators are built from two cytoplasmic membrane proteins, which for simplicity are generically referred to here as MotA and MotB. MotA is predicted to contain four transmembrane helices (TMH) with a large cytoplasmic insertion between TMH2 and TMH3. MotB is predicted to contain a short cytoplasmic sequence, a single TMH, and a C-terminal peptidoglycan binding (PGB) domain. Early biochemical work defined the stator stoichiometry as MotA_4_B_2_^11^, and this subunit composition has informed attempts to derive mechanism for conversion of ion flow into rotation (reviewed in ^12^). Extensive experimental studies have led to a model of stator function in which docking of the MotA cytoplasmic loop to the rotor C-ring simultaneously induces ion permeation through the stator and release of the MotB-PGB domain to bind to the peptidoglycan (PG) surrounding the flagellar basal body^13,14^. Ion flow is proposed to lead to conformational changes in the cytoplasmic domain of MotA that generate torque in the rotor^15–17^. In the absence of a stator structure various mechanistic hypotheses have been proposed to explain the coupling of ion flow to conformational change, most of which explicitly use the predicted 2-fold symmetry of a MotA_4_B_2_ complex^17–19^.

### Flagellar stators are MotA_5_B_2_ complexes

We used cryo-electron microscopy (cryo-EM) to study stator complexes from a range of bacterial species with different ion specificities (Extended Data Fig. 1). Two-dimensional class averages of the complexes from three species (*Vibrio mimicus, Clostridium sporogenes* and *Bacillus subtilis*) clearly showed a distorted, pentagonal, structure (Fig. 1b). 3D reconstructions of these complexes yielded volumes that could only be interpreted as MotA_5_B_2_ assemblies (Fig. 1c; Extended Data Fig. 2), with five copies of MotA fully enclosing the TMHs of two copies of MotB (Fig. 2a). Although we do not observe the PGB-domains of MotB in the resolved structures, these domains must be present in the imaged complexes because the stators were purified using an affinity tag located after the PGB domain. Thus, the PGB-domain of MotB has no fixed location with respect to the core complex in the context of the isolated protein. The stator structures are compatible with sequence conservation data, with inter- and intra-molecular co-evolution data, and with published cysteine crosslinking^20,21^ and tryptophan scanning mutagenesis^22,23^ (Extended data Fig. 3).

**Figure 2.**
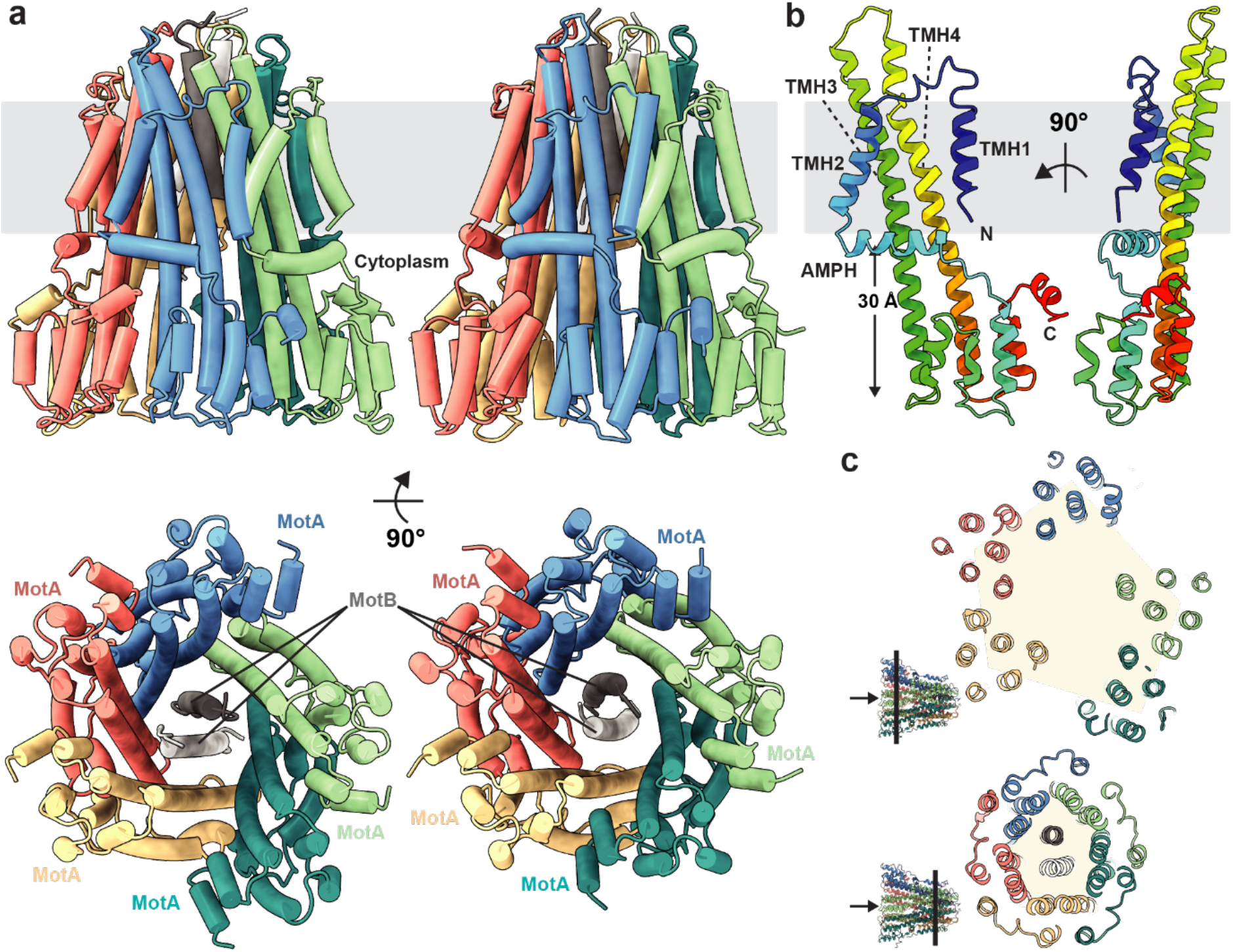
Structures of stators from *C. sporogenes* and *B. subtilis*. **a**, The *C. sporogenes* (left) and *B. subtilis* (right) stators are shown as cartoon representations and coloured as in Fig. 1c. Upper panel, side view with membrane indicated in grey. Lower panel, view from the cytoplasm. **b**, Two views of a single MotA subunit (*C. sporogenes*) coloured from blue at the N-terminus to red at the C-terminus. **c**, Slabs (viewed from cytoplasm) through the *C. sporogenes* complex at the indicated positions on the inset structure (arrow indicates the cytoplasmic side of the complex). Distortion of the MotA subunits from a regular pentagon arrangement becomesmore extreme in the cytoplasmic regions.

The four TMHs of MotA are arranged in two layers. TMH3 and TMH4 line the central pore, while TMH1 and TMH2 form a surrounding outer layer of helices (Fig. 2a,b). TMH1 and TMH2 are not in contact with each other within a single subunit but instead interact between adjacent subunits, thereby stabilising the MotA assembly. Immediately following TMH2 there is an amphipathic helix (AMPH) running perpendicular to the TMHs at the cytoplasmic membrane surface, with the five copies of this helix forming a belt around the outside of the structure. TMH3 and TMH4 extend 30 Å outside the membrane to form the core of the MotA cytoplasmic domain, with the rest of the domain built from helices inserted in the loop between the AMPH and TMH3. Both within and outside the membrane domain the pentameric arrangement of MotA is distorted (Fig. 2c). Charged residues shown to be essential for the interaction of the stator with the rotor C-ring^24^ are located towards the base of this domain, forming a ring that decorates the surface of the pentamer (Extended data Fig. 4).

The TMHs of the two copies of MotB are located in the central pore of the distorted MotA pentamer, with their hydrophobic sidechains completely buried within the MotA ring. From the N-terminal ends of the MotB TMHs clear densities extend down to contact the inner surfaces of TMH3 and TMH4 in the cytoplasmic domains of MotA (Fig. 3a,b; Extended Data Fig. 5a,b). Although the densities are too weak for the sequence to be traced, they are of sufficient length to account for most of the MotB N-terminus, including a cluster of positive charges essential for motor function^25^. At the non-cytoplasmic face of the complex the MotB TMHs emerge vertically from the MotA pentamer and are followed by another short helical section that packs down between the TMH3-TMH4 loops of the MotA chains. The connectivity of these densities defines them as the plug helices previously implicated by mutagenesis as critical to sealing the complexes in an off state^26^.

**Figure 3.**
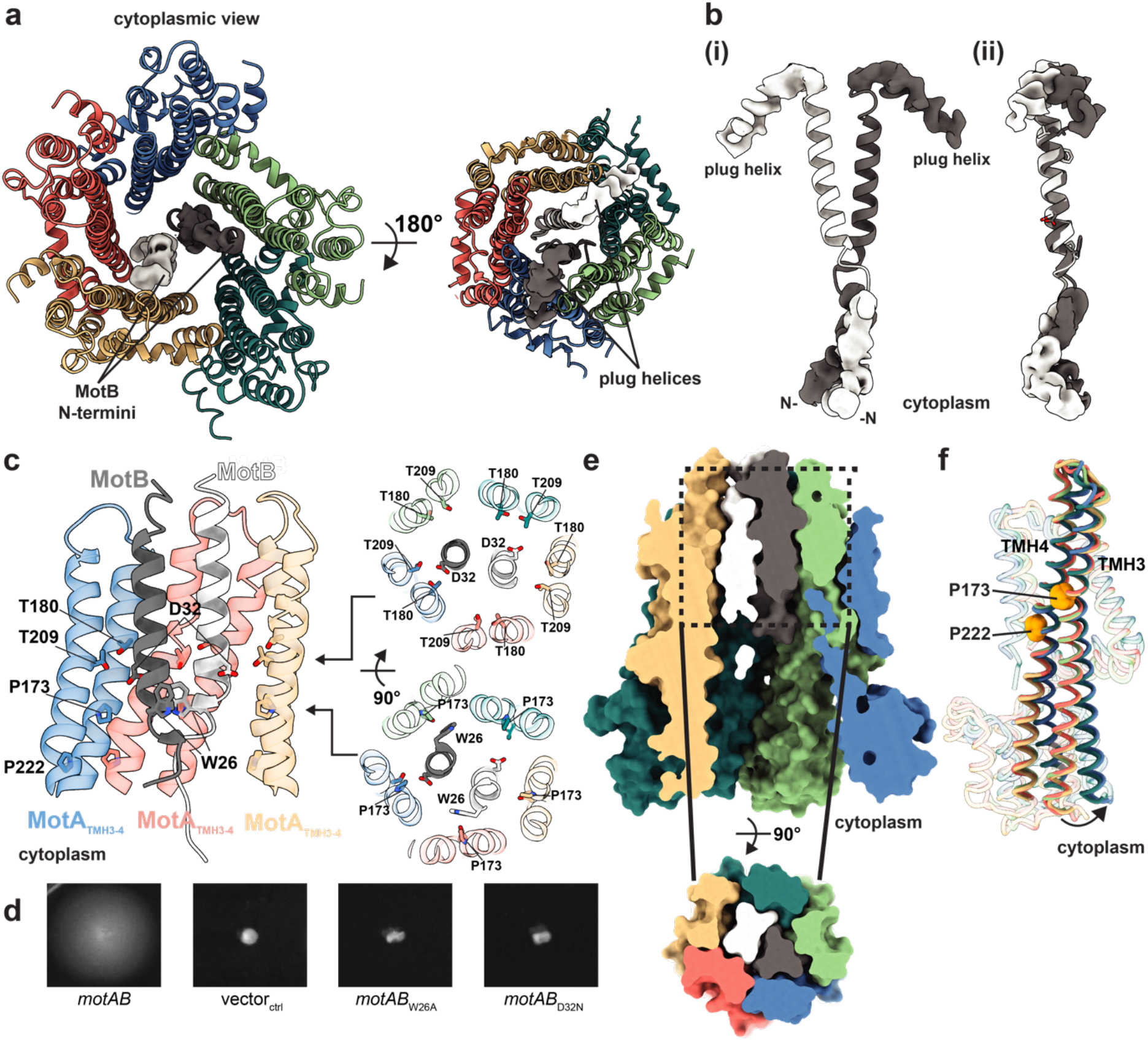
Functionally critical regions of the stator complex. **a**, Structure of the *C. sporogenes* stator shown as cartoon representations coloured as in Figs. 1 and 2 with the unmodeled density for the MotB N-terminal extensions (Left) and plug helices (Right) shown. **b**, (i) Isolated MotB dimer extracted from the *C. sporogenes* stator and (ii) superposition of the TMHs of the two MotB chains showing the relative rotation of the N-terminal extensions and plug helices. **c**, The environment around MotBDp32 within the membrane. (Left) Only the core MotA helices within the transmembrane region are shown and the two copies of MotA at the front of the view are removed. (Right) Slabs through the stator core at the indicated heights. Residue numbering is that of the *E. coli* MotAB stator but displayed on the *C. sporogenes* stator structure. **d**, Motility in soft agar of *E. coli* RP6894 (Δ*motAB*) complemented with plasmids expressing *motAB* with the indicated mutations or the vector control (vectorctrl). **e**, Surface representation of the model shows close packing. (Top) Side view with front of complex removed. (Bottom) Top-down view of the slab indicated by dashed lines. **f,** Overlay of the five copies of the *C. sporogenes* MotA chain reveals they fall into two conformational classes which differ in the degree of flexing at the highlighted prolines.

Prior mutagenesis studies have established that a series of conserved residues in the TMHs of MotA and MotB are important for flagellar motion and/or ion-flow through the stator (reviewed in^3^). Invariant MotB_D32_ (using the *Escherichia coli* numbering system) is the key protonatable residue and both copies are seen to lie within a ring formed by the five copies of another invariant polar residue, MotA_T209_ (Fig. 3c). A second Thr residue (at a position corresponding to residue A180 in *E. coli* MotA) that is conserved in the Na^+^-dependent stators also contributes to this ring, and forms part of a track of Na^+^/H^+^ specificity determining residues that line the inner surface of the MotA pore (Extended data Fig. 1b). Two conserved Pro residues in MotA have been shown to be important for torque generation^27^. One of these, MotA_P222_ can now be seen to be required for contacts between neighbouring MotA monomers. The other, MotA_P173_, forms a second ring of conserved residues with invariant MotA_Y217_, two helical turns down from the Thr ring. This hydrophobic ring contacts MotB at the completely conserved MotB_W26_. A MotB_W26A_ substitution completely abolished motility confirming the importance of this contact (Fig. 3d).

### Asymmetry and the implications for activation of ion flow

The 5:2 stoichiometry of the stator complex leads to multiple levels of asymmetry in the structure (Fig. 2c; Extended Data Fig. 5c). The pentagon formed by the MotA subunits within the membrane is distorted to accommodate and seal around the two MotB TMHs. The asymmetry of this part of the complex is also driven by the two MotB plug helices sitting between the MotA loops, which divide the MotA chains into two groups separated in the extracytoplasmic region. Removal of the MotB plug has been shown to lead to uncontrolled ion flow through the MotAB channel^26^. However, our structures show that the plug helices are not the sole block to ion permeation since there are no detectable channels across the cytoplasmic membrane compartment (Fig. 3e; Extended Data Fig. 5d). Embedding plug-free structures in full lipid bilayer models and running extended simulations demonstrated the observed structures are stable, low energy, states (Extended Data Fig. 6). No ion permeation across the bilayer was seen in any simulation, supporting the idea that the complexes currently seen will require rearrangement for activity.

Activation of ion flow is proposed to be triggered by docking of the inactive stator onto the flagellar C-ring via the MotA cytoplasmic domains resulting in signal propagation from the cytoplasm to the plug region and plug release^13,14^. Our structures reveal two potential routes for such a signal. The first involves the cytoplasmic N-termini of the MotB subunits which contain functionally essential residues^25^ that interact with the inside of the MotA pentamer through highly evolutionarily coupled contacts (Extended data Fig. 3d). C-ring-induced movement of MotA would be communicated to MotB at this site leading to alterations at the opposite end of the MotB TMHs. The second possible route of signal propagation is directly through the MotA subunits, with hinging of the long TMH3 and TMH4 helices altering the conformation of the plug helix binding loops to allow plug release. Our structures provide insight into the conformational changes that the MotA cytoplasmic domains can undergo. The structures show differing degrees of hinging of the MotA cytoplasmic domains relative to the membrane embedded helices (Fig. 3f; Extended Data Fig. 5e) suggesting changes in the degree of asymmetry may link to functional state. Our structures also reveal that the two MotA residues known to be essential for interaction with the C-ring protein FliG^24^ are located on opposite sides of the MotA cytoplasmic domain, with MotA_R90_ from one copy facing MotA_F98_ from the neighbouring copy (Extended data Fig. 4). Therefore docking of the C-terminal domain of FliG between two MotA subunits could trigger conformational change in the stator.

### Coupling of ion-flow to flagellar rotation

The most striking feature of the asymmetry of the 5:2 subunit stoichiometry is that it places the TMHs of the two copies of MotB, including the critical MotB_D32_ residue, in different environments within the distorted MotA pentagon. The system is therefore primed for differential binding of H^+^ or Na^+^ at the critical MotB_D32_ residue to induce changes in the relative positioning of the MotB and MotA helices, most likely by rotation of one relative to the other. Because MotB becomes tethered to the PG upon stator activation^13^ (Fig. 4a) it follows that the MotA ring rotates around the MotB dimer. The 5:2 subunit stoichiometry implies a model whereby a single channel opens to allow ion binding to MotB_D32_ on one MotB triggering rotation of the MotA ring by ~36° (Fig. 4b). This motion would bring the second MotB chain into the same position relative to the surrounding MotA subunits as the starting arrangement of the first MotB chain, closing the first channel and opening the second. Each subsequent ion binding event would trigger a further ratchet motion of 36°, with each turn of the MotA cytoplasmic domains providing a “power stroke” to the rotor. An alternative model in which the second ion binding event leads to a reset to the original position and full rotation does not occur, would also be compatible with the structure. Such a mechanism would still be capable of driving full rotation of the rotor component, acting like an energised escapement mechanism.

**Figure 4.**
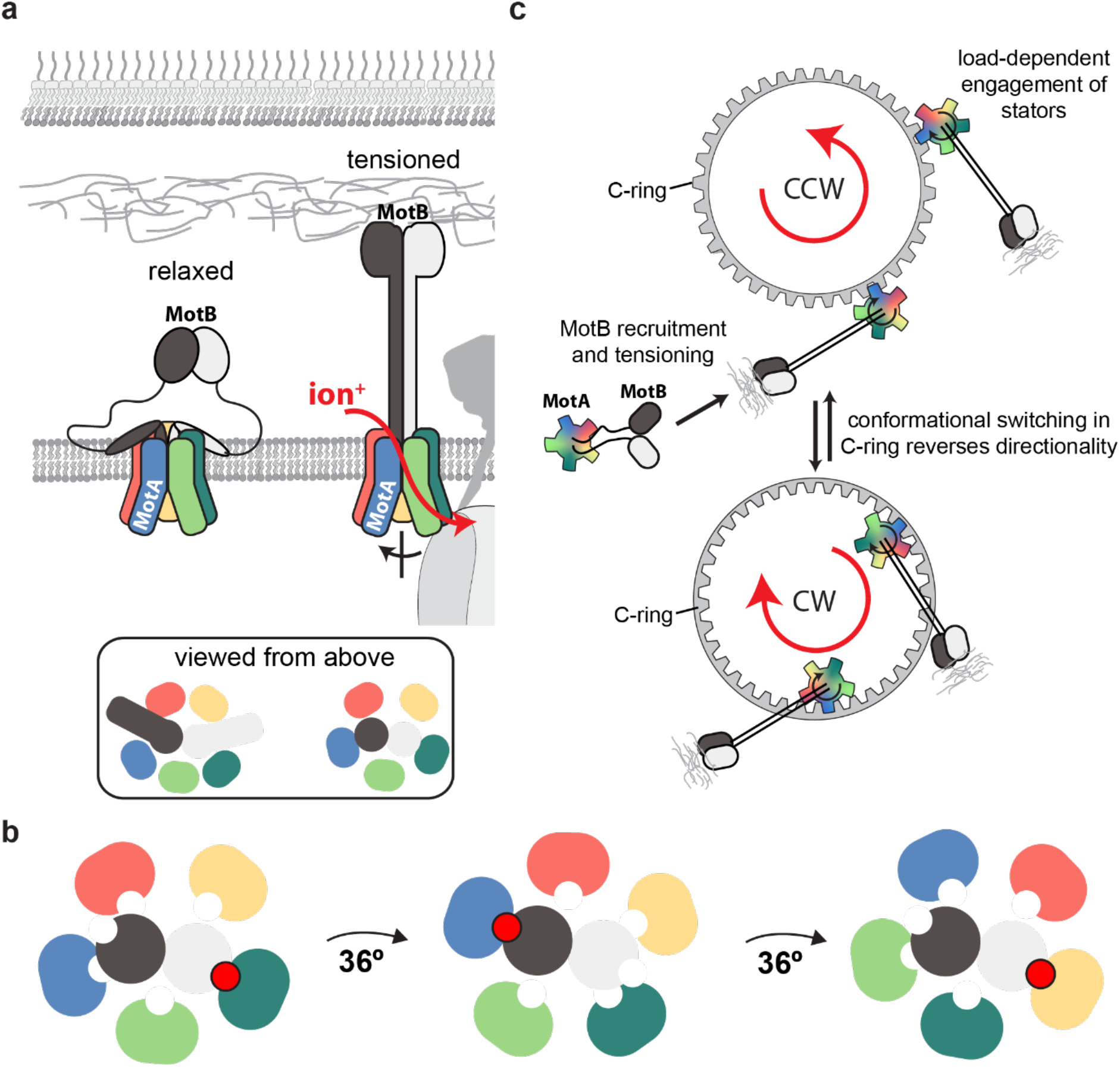
Mechanistic model for the generation of bi-directional flagellar torque. **a**, Activation of the stator complex from the structurally resolved state (here termed ‘relaxed’) to form a ‘tensioned’ state permissive to ion-flow. This conformational change is likely driven by interactions between the C-terminal peptidoglycan binding domains (black and white ovals) of MotB and the peptidoglycan layer, as well as interactions between the stator and the C-ring complex. **b**, Cartoon showing top views of the intra-membrane core of the stator complex with five MotA chains surrounding two MotB subunits. Bound ions are shown as red spheres. Ion flow leads to rotation of the MotA ring through alternating formation of MotB-ion-MotA interactions by the two MotB chains that processes around the surrounding MotA subunits. **c**, Model describing how a stator that rotates in one direction can drive either clockwise (CW) or counterclockwise (CCW) rotation of the flagellum depending on the conformational state of the C-ring.

Any mechanism for coupling ion flow to flagellar rotation must also explain how the direction of rotation of the flagellum can reverse in response to chemotactic stimuli. All experimental evidence (reviewed in ^28^) shows that the chemotaxis machinery acts on the FliG subunit of the C-ring rather than the stator. Our unidirectional rotation model for the stator mechanism can account for flagellar reversal if the chemotaxis-linked conformational changes induced in the C-ring lead to an alteration in the side of the stator that is driving the rotation (Fig. 4c). Consistent with this proposal large conformational changes in the stator-interacting FliG component of the C-ring have been observed in crystal structures of FliG fragments, where 180° rotations of the C-terminal domain relative to the middle domain have been observed^29^. The model also predicts that any reversal of the ion flow through the stator would have the potential to reverse the direction of flagellar rotation even in the absence of switching by the chemotaxis machinery, and this phenomenon has been observed in *Streptococcus* species assayed under high pH conditions^30,31^.

### Common architecture across multiple bacterial ion-driven machines

The MotAB system is related at the sequence level to the ExbBD complex found in Gram-negative bacteria that uses ion-flow across the cytoplasmic membrane to power transport processes at the outer membrane via the trans-periplasmic TonB protein^32^. We determined cryo-EM structures of ExbBD complexes from *E. coli* and *Pseudomonas savastanoi* (Extended Data Fig. 7). Both displayed a 5:2 ExbB:ExbD stoichiometry that differs from the subunit composition of earlier structures^33,34^, but agrees with the subunit stoichiometry of a novel structure of the *E. coli* ExbBD reported whilst this manuscript was in preparation^35^. Comparison of these new ExbB5D2 structures to the stator complexes reveals a high level of structural conservation, particularly within the membrane domain (Extended Data Fig. 8a,b). Both the flattened pentagon geometry and the alignment of mechanistically important residues, such as the conserved Asp within a ring of Thr residues, suggest that the two systems use the same molecular mechanism. We therefore predict that the ExbB will rotate relative to the ExbD helices in response to proton flow. Outside the core TMH region there are structural differences between the systems that presumably reflect their very different biologies. ExbB is very differently elaborated relative to MotA, with only one TMH packing across the pair of helices that form the core inner ring and no bracing helices strengthening packing between subunits (Extended Data Fig. 8c). The ExbB cytoplasmic domains are only superficially related to the corresponding MotA domain and lack the short pair of C-terminal helices found in MotA (Extended Data Fig. 8d).

*P. savastanoi* ExbB and ExbD were purified as a complex with TonB when all three proteins were co-expressed (Extended data Fig. 9a). However, no extra density was observed in the cryo-EM maps of this complex relative to the ExbBD complex alone, suggesting that TonB is located on the outside of the ExbBD complex and dissociates upon sample freezing. A peripheral location for TonB is consistent with both co-evolution and mutagenesis/suppressor data^36^, which suggest that the TonB binding site is on the outside of the ExbB transmembrane domain (Extended data Fig. 9b). TonB consists of a single pass TMH, followed by an extended periplasmic region that interacts with the periplasmic domain of ExbD^37^, and terminates in a folded domain that links with outer-membrane receptor proteins^38^. We speculate that the TonB TMH packs against the exterior of the ExbBD complex so that conformational change in TonB is driven by rotation of the ExbB component relative to ExbD. By extension, the homologous TolQRA system will also share this architecture and be mechanistically related^39^.

Bacteria from the *Bacteroidetes* phylum possess a motor complex that harvests energy from ion-flow to drive protein secretion and to power bacterial motility via a non-flagellar mechanism termed gliding motility^40^. The structure of this complex is described in a companion paper (Hennell-James et al, companion paper). Although the constituent GldL and GldM subunits of this motor have no sequence similarity to the subunits of the MotAB or ExbBD complexes, the *Bacteriodetes* motor complex exhibits the same 5:2 subunit stoichiometry as these complexes (Fig. 5a). All three complexes have an intramembrane core consisting of a central subunit TMH dimer surrounded by a 10 TMH ring. Structural comparisons demonstrate the similarity between the three motors in the arrangement of this intramembrane core and of the height within the membrane at which charged residues critical to function are located (Fig. 5b,c). Such shared underlying architecture between otherwise highly dissimilar motors (Fig. 5d) implies an unexpected commonality in their mechanism.

**Figure 5.**
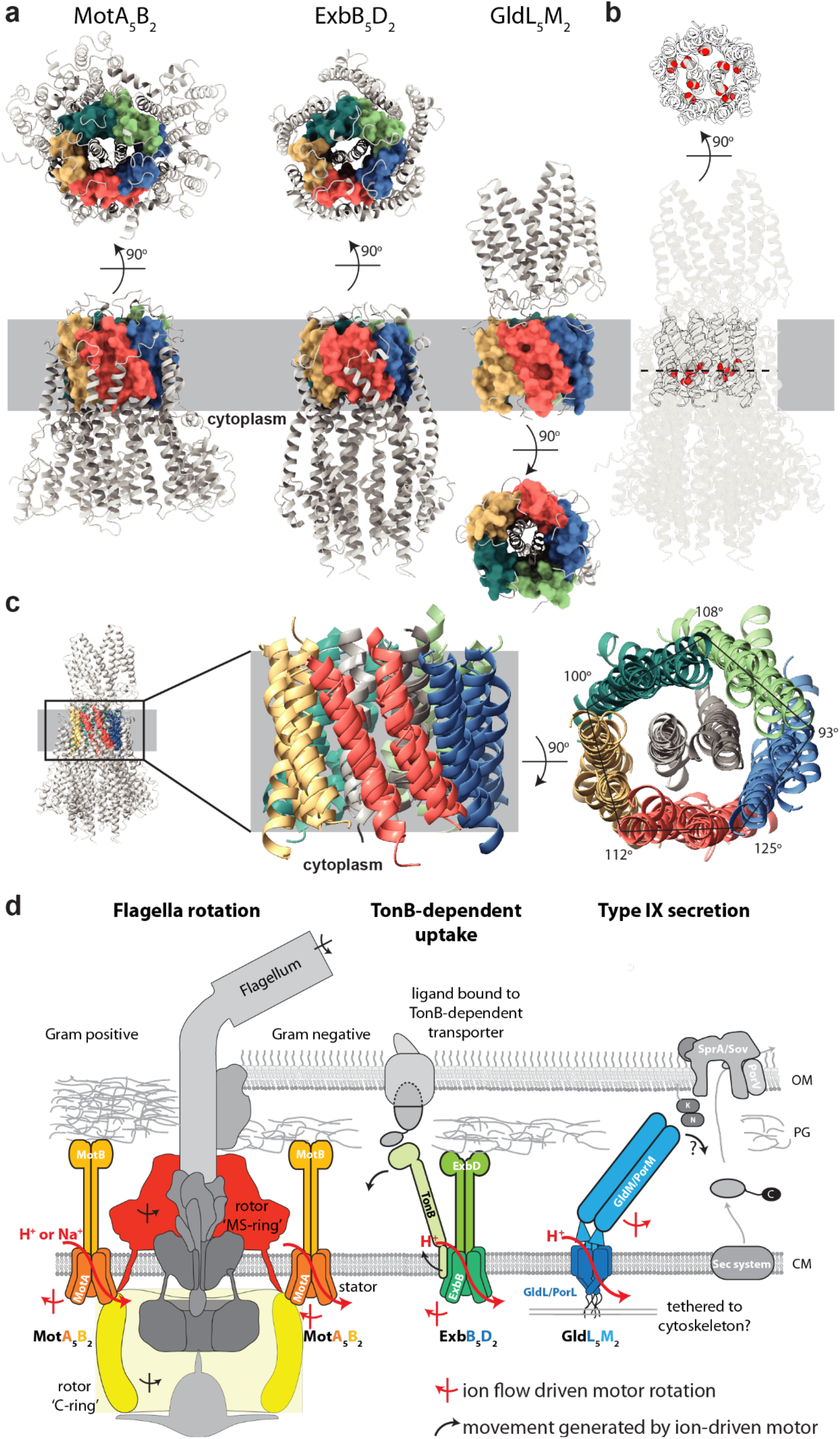
Conservation of core architecture between diverse families of ion-driven motors. **a**, Representatives of three ion-driven motor families that share a common structural core. Complexes are shown as grey cartoons with helices equivalent to those in the MotA inner ring displayed in coloured surface representation. **b**, Overlay of the three complexes (common core in grey cartoons, other structure semi-transparent cartoons). Mechanistically essential charged residues within the common core (space filling side chains; C, grey; O, red) occur at the same height with respect to the membrane irrespective of whether they occur on the MotA- or MotB-equivalent chain. **c**, Overlay of the common core of the three complexes. The distortion from pentamer symmetry within the membrane is shared between all three families **d**, Cartoon summarising the updated view of how the three families of ion-driven motors are coupled to their different biological effects. Note that ion movement drives rotation of the central subunits in GldLM but of the peripheral subunits in MotAB/ExbBD. OM, outer membrane; CM, cytoplasmic membrane; PG, peptidoglycan

## Acknowledgements

We thank David Blair for providing *E. coli* RP6894. We thank E. Johnson and A. Costin of the Central Oxford Structural Molecular Imaging Centre (COSMIC) for assistance with data collection, and H. Elmlund (Monash) for access to SIMPLE code ahead of release. We acknowledge the use of the Central Oxford Structural Microscopy and Imaging Centre (COSMIC). The Central Oxford Structural Microscopy and Imaging Centre is supported by the Wellcome Trust (grant no. 201536), The EPA Cephalosporin Trust, The Wolfson Foundation and a Royal Society/Wolfson Foundation Laboratory Refurbishment Grant (no. WL160052). Research in S.M.L.’s laboratory is supported by a Wellcome Trust Investigator Award (grant no. 100298), a Collaborative award (no. 209194) and an Medical Research Council (London) Programme Grant (no. MR/M011984/1). Research in B.C.B.’s laboratory is supported by a Wellcome Trust Investigator Award (grant no. 107929/Z/15/Z). Research in J.W.C.’s laboratory is supported by the Canadian Institutes of Health Research (grant 178048-BMA-CFAA-11449). Research in P.J.S.’s lab is funded by Wellcome (208361/Z/17/Z), the MRC (MR/S009213/1) and BBSRC (BB/P01948X/1, BB/R002517/1 and BB/S003339/1). This project made use of time on ARCHER and JADE granted via the UK High-End Computing Consortium for Biomolecular Simulation, HECBioSim (http://hecbiosim.ac.uk), supported by EPSRC (grant no. EP/R029407/1), and Athena at HPC Midlands+, which was funded by the EPSRC on grant EP/P020232/1, and used the University of Warwick Scientific Computing Research Technology Platform for computational access.

## Author contributions

J.C.D carried out all biochemical work except as credited otherwise, prepared cryo-EM grids, collected and processed EM data and determined the structures. J.C.D., S.J., and S.M.L designed the project, interpreted the data, built models, and wrote the first draft of the paper. S.J. also performed MALS experiments. O.V. and P.J.S. performed molecular dynamics simulations. A.M., H.M., and T.G. carried out biochemical work on *Pseudomonas* TonB-ExbB-ExbD. R.H.J. and B.C.B. contributed the GldLM structure. J.W.C. initiated and provided materials for the ExbBD project. All authors commented on drafts of the manuscript.

## Methods

### Bacterial strains and plasmids

Bacterial strains and plasmids used in this study are listed in Supplementary Table 1. The pT12 backbone used for all protein expression was derived from Kuhlen *et al* ^41^. Plasmids were generated by Gibson assembly of PCR fragments using the NEBuilder HiFi Master Mix (NEB). Fragments were created by PCR with the relevant primers (listed in Supplementary Table 2) using Q5 polymerase (NEB) and genomic DNA templates obtained from the Liebniz Institute [dsmz.de]: *Vibrio mimicus* (DSM 19130), *Bacillus subtilis* 168 (DSM 402), *Clostridium sporogenes* 388 (DSM 795), *Escherichia coli* W (DSM 1116), *Pseudomonas savastanoi*, pv. phaseolicola 1448A (DSM 21482). Gibson assembly and PCR were carried out following the manufacturer’s recommendations. *E. coli* RP6894 (Δ*motAB*) for motility assays was generated by J. S. Parkinson and gifted by D.F. Blair.

### Purification of MotAB/PomAB and ExbBD complexes

*V. mimicus* PomAB, its derivative PomAB_Δ61-120_ lacking unstructured periplasmic residues of PomB, *B. subtilis* MotAB, *C. sporogenes* MotAB, *E. coli* ExbBD, and *P. savastanoi* TonB-ExbBD complexes were expressed in *E. coli* MT56 as a single operon from a pT12 vector encoding a C-terminal twin-strep tag. Purification steps were similar across all constructs and carried out at 4 °C. Briefly, cells were grown at 37 °C for 16 h in TB media containing kanamycin (50 μg/mL) and rhamnose monohydrate (0.1% w/v) then collected by centrifugation at 4,000*g*. Cell pellets were resuspended in TBS (100 mM Tris, 150 mM NaCl, 1 mM EDTA pH 8.0) plus 30 μg/mL DNase I and 400 μg/mL lysozyme for 30 mins before passage through an EmulsiFlex C5 homogenizer (Avestin) at 15,000 psi. Unbroken cells were removed by centrifugation at 24,000*g* for 20 min. The supernatant was recovered and total membranes were collected by centrifugation at 200,000*g* for 1.5 h. Membranes were resuspended in TBS and solubilized by incubation with 1% (w/v) lauryl maltose neopentyl glycol (LMNG; Anatrace) for 2 h. Insoluble material was removed by centrifugation at 100,000*g* for 30 min. Solubilized membranes were then applied to a Streptactin XT column (IBA Lifesciences). The resin was washed with 10 column volumes (CV) of TBS containing 0.02% (w/v) LMNG and proteins were eluted in 5 CV of TBS supplemented with 0.01% (w/v) LMNG and 50 mM D-biotin (IBA Lifesciences). Eluates were concentrated using a 100-kDa molecular weight cutoff (MWCO) Vivaspin 6 (GE Healthcare) centrifugal filter unit and injected onto a Superose 6 Increase 10/300 GL size exclusion column (GE Healthcare) pre-equilibrated in TBS plus 0.01% (w/v) LMNG. Peak fractions were collected and concentrated using a 100-kDa MWCO Vivaspin 500 (GE Healthcare) centrifugal filter unit.

For *P. savastanoi* TonB-ExbBD and ExbBD complexes, SEC-MALS analysis was carried out by injecting 100 μL (A_280nm_ = 1.0) of either sample onto a Superose 6 increase 10/300 GL (GE Healthcare) equilibrated in TBS containing 0.02% (w/v) LMNG. Light scattering and refractive index changes were measured using a Dawn Heleos-II light-scattering detector and an Optilab-TrEX refractive index monitor. Analysis was carried out using ASTRA 6.1.1.17 software using a theoretical extinction coefficient of 1.02 (Abs_0.1%_) and a protein dn/dc value of 0.186 mL/g and a detergent dn/dc value of 0.143 mL/g.

### Cryo-EM sample preparation and imaging

Purified complexes (4 μL each) of *V. mimicus* PomAB (A_280nm_ = 0.5), PomAB_Δ61-120_ (A_280nm_ = 0.55), *B. subtilis* MotAB (A_280nm_ = 1.0), *C. sporogenes* MotAB (A_280nm_ = 0.8), *E. coli* ExbBD (A_280nm_ = 3.2), or *P. savastanoi* TonB-ExbBD (A_280nm_ = 2.0) were adsorbed to glow-discharged holey carbon-coated grids (Quantifoil 300 mesh, Au R1.2/1.3) for 10 s. Grids were then blotted for 2 s at 100% humidity at 8°C and frozen in liquid ethane using a Vitrobot Mark IV (FEI). Alternatively, specimens were prepared by supplementing *V. mimicus* PomAB (A_280nm_ = 2.3), PomAB_Δ61-120_ (A_280nm_ = 3.7), *B. subtilis* MotAB (A_280nm_ = 7.2), *C. sporogenes* MotAB (A_280nm_ = 8.6), *E. coli* ExbBD (A_280nm_ = 4.2) with 0.7 mM fluorinated octyl maltoside (fluo OM; Anatrace) prior to grid preparation.

Data were collected in counting mode on a Titan Krios G3 (FEI) operating at 300 kV with a GIF energy filter (Gatan) and K2 Summit detector (Gatan) using a pixel size of 0.822 Å and a total dose of 48 e^-^/Å^2^ spread across 20 or 32 fractions. Except for *P. savastanoi* TonB-ExbBD, all datasets included movies from grids prepared with and without the presence of fluo OM to improve distribution of particle orientations.

### Cryo-EM data processing

Motion correction and dose weighting were performed using MotionCor implemented in Relion 3.0^42^. Contrast transfer functions were calculated using CTFFIND4^43^. Particles were picked in Simple^44^ and processed in Relion 3.0^42^. Gold standard Fourier shell correlations using the 0.143 criterion and local resolution estimations were calculated within Relion^42^ (Extended Data Fig. 2).

*V. mimicus* PomAB particles (1,172,445) underwent one round of reference-free 2D classification, from which 253,681 particles were selected and used to generate an *ab initio* initial model. This model was low-pass filtered to 30 Å and used as reference for 3D classification, generating a class that refined to 6.8 Å from 155,280 particles.

For the deletion construct PomAB_Δ61-120_ that improved particle orientations and data quality, particles (2,383,062) were extracted from 13,980 movies. Following one round of reference-free 2D classification, 800,844 particles were classified in 3D (4 classes) against a 40 Å low-pass filtered map of PomAB. A class containing 244,654 particles was further subjected to masked refinement yielding a 4.8 Å map. Refinement after Bayesian particle polishing and per-particle defocus with beamtilt estimation further improved map quality to 4.2 Å.

*B. subtilis* MotAB particles (1,532,430) were extracted over 11,588 movies. After 2D classification, selected particles (397,584) underwent two rounds of 3D classification (3 classes each) using a 40 Å low-pass filtered map generated from a subset of particles refined against a 60 Å low-pass filtered map of PomAB_Δ61-120_. A class made up of 122,615 particles was refined to 3.9 Å. Bayesian particle polishing further improved map resolution by 0.2 Å, and subsequent CTF refinement using per-particle defocus with beamtilt estimation generated a 3.5 Å map. To improve MotB N-terminal and plug densities, a subset of fluorinated particles (43,375) was selected and refined against the 3.5 Å reconstruction, generating a 5.0 Å map that was used to depict these regions in Extended Data Fig. 5.

*C. sporogenes* MotAB particles (1,998,900) were extracted from 9,148 movies and subjected to a round of reference-free 2D classification. Initial 3D classification performed against a 60 Å low-pass filtered map of *B. subtilis* MotAB revealed two prominent classes which represented a monomeric MotAB complex and a non-physiological end-to-end dimer of MotAB. These classes were used as references in a supervised multi-reference 3D classification against the full 1,137,357 particle set to exclude dimeric particles. Unsupervised 3D classification (4 classes) performed against 865,446 monomeric particles and further refinement yielded 3.8 Å from 314,230 particles. Bayesian particle polishing followed by per-particle defocus with beamtilt estimation further improved map quality to 3.4 Å.

Movies (6,902) were collected for *E. coli* ExbBD, resulting in the extraction of 2,045,350 particles. Following one round of reference-free 2D classification, an initial model of ExbBD was generated by 3D classification and refinement of a particle subset against a 40 Å low-pass filtered 5:1 ExbBD complex^34^ (EMD-6928). The resulting map was used as initial model for multiple rounds of 3D classification against the full 2D-classified particle set (755,677). After refinement of 227,700 particles, this protocol generated a 5.8 Å map, improving to 4.6 Å following Bayesian particle polishing and per-particle defocus plus beamtilt estimation.

Movies (4,232) were collected for *P. savastanoi* TonB-ExbBD and 1,342,900 particles were extracted. Particles were subjected to two rounds of 2D classification with centered re-extraction between classifications. The cleaned 499,697 particles were subjected to C5-symmetric 3D classification against a 40 Å low-pass filtered 5:2 ExbBD map from *E. coli*. The resultant 202,356 particles were refined with C5 symmetry to generate a 3.5 Å map that lacked density for the transmembrane helices (TMHs) of ExbD. Particles were polished and subjected to an additional round of 2D classification followed by 3D classification with C1 symmetry, resulting in 3.9 Å map from 110,164 particles after refinement in C1. An additional round of Bayesian polishing and refinement (C1) yielded a 3.8 Å map. Per-particle defocus and beamtilt estimation followed by alignment-free 3D classification and subsequent local refinement (C1) yielded a 3.8 Å map with improved density for the TMHs of ExbD from 65,617 particles.

### Model building and refinement

Atomic models were built using Coot^45^. Multiple rounds of rebuilding (in both the globally sharpened and local-resolution filtered maps) and real-space refinement in Phenix^46^ using secondary structure, rotamer and Ramachandran restraints yielded the final models described in Table 1. All models were validated using Molprobity^47^. Conservation analysis was carried out using the Consurf server^48^. A homology model of *E. coli* MotAB was generated by sequence threading against the *Clostridium* model using Phyre2^49^. Figures were prepared using UCSF ChimeraX^50^ and Pymol (The PyMOL Molecular Graphics System, Version 2.0 Schrödinger, LLC). All models depicted in figures are based on the highest resolution *Clostridium* model, unless otherwise specified. Residue numbering adopts the reference *E. coli* sequence and model; a residue conversion table is provided (Supplementary Table 3).

**Table 1.** Cryo-EM data collection, refinement and validation statistics.

|  | *C. sporogenes*<br>MotAB<br>(EMDB-10895)<br>(PDB ID 6YSF) | *B. subtilis*<br>MotAB<br>(EMDB-10899)<br>(PDB ID 6YSL) | *V. mimicus*<br>PomAB<br>(EMDB-10901) | *E. coli*<br>ExbBD<br>(EMDB-10902) | *P. savastanoi*<br>ExbBD<br>(EMDB-10897) |
| --- | --- | --- | --- | --- | --- |
| **Data collection and processing** |  |  |  |  |  |
| Magnification | 165,000 | 165,000 | 165,000 | 165,000 | 165,000 |
| Voltage (kV) | 300 | 300 | 300 | 300 | 300 |
| Electron exposure (e-/Å<sup>2</sup>) | 48 | 48 | 48 | 48 | 48 |
| Defocus range (µm) | 1.0-3.0 | 1.0-3.0 | 1.0-3.0 | 1.0-2.5 | 1.0-3.0 |
| Pixel size (Å) | 0.822 | 0.822 | 0.822 | 0.822 | 0.822 |
| Symmetry imposed | C1 | C1 | C1 | C1 | C1 |
| Initial particle images (no.) | 1,998,900 | 1,532,430 | 2,383,022 | 2,045,350 | 1,342,937 |
| Final particle images (no.) | 314,230 | 122,615 | 244,654 | 227,700 | 65,617 |
| Map resolution (Å) | 3.4 | 3.5 | 4.2 | 4.6 | 3.8 |
| FSC threshold | 0.143 | 0.143 | 0.143 | 0.143 | 0.143 |
| Map resolution range (Å) | 3.2-5.2 | 3.3-6.0 | 3.9-6.5 | 4.0-6.5 | 3.6-6.1 |
| **Refinement** |  |  |  |  |  |
| Initial model used (PDB code) | None | None |  |  |  |
| Model resolution (Å) | 3.4 | 3.5 |  |  |  |
| FSC threshold | 0.143 | 0.143 |  |  |  |
| Model resolution range (Å) | 3.2-5.2 | 3.3-6.0 |  |  |  |
| Map sharpening *B* factor (Å<sup>2</sup>) | -117 | -104 |  |  |  |
| Model composition |  |  |  |  |  |
| Non-hydrogen atoms | 10220 | 10128 |  |  |  |
| Protein residues | 1327 | 1324 |  |  |  |
| Ligands | 0 | 0 |  |  |  |
| *B* factors (Å<sup>2</sup>) |  |  |  |  |  |
| Protein | 53 | 99 |  |  |  |
| Ligand | NA | NA |  |  |  |
| R.m.s. deviations |  |  |  |  |  |
| Bond lengths (Å) | 0.004 | 0.007 |  |  |  |
| Bond angles (°) | 0.757 | 1.425 |  |  |  |
| Validation |  |  |  |  |  |
| MolProbity score | 2.15 | 1.85 |  |  |  |
| Clashscore | 13.66 | 5.93 |  |  |  |
| Poor rotamers (%) | 0.90 | 0.47 |  |  |  |
| Ramachandran plot |  |  |  |  |  |
| Favored (%) | 91.47 | 90.92 |  |  |  |
| Allowed (%) | 8.53 | 9.01 |  |  |  |
| Disallowed (%) | 0.00 | 0.08 |  |  |  |

### Evolutionary covariance analysis

Coevolutionary contacts for *E. coli W* MotA were determined by the Gremlin web-server^5^. Searches used the Jackhmmer algorithm for multiple sequence alignment, an E-value threshold of 10^-10^ and a minimum coverage of 75%. Intra- and intermolecular contacts were mapped to the *E. coli* MotA structure using Gremlin beta^5^. Intermolecular contacts between MotA and MotB (residues 1-120) were determined using an E-value threshold of 10^-20^ and 10^-2^, respectively. Intermolecular contacts between TonB and ExbB were determined using an E-value threshold of 10^-20^. Contacts with a probability score greater than 0.9 were regarded as significant and listed in Supplementary Table 4.

### Simulation setup

All simulations were run using GROMACS 2018^51^. The systems were initially setup using the Martini 2.2 coarse-grain (CG) force field and solvated with water and 0.15 M NaCl to neutralise the system^52^. The membranes were constructed using INSANE with a 4:1 ratio of POPE:POPG lipids^53^. An elastic network of 1000 kJ mol^-1^ nm^-2^ was applied between all backbone beads between 0.5 and 1 nm. Electrostatics were described using the reaction field method, with a cut-off of 1.1 nm using the potential shift modifier and the van der Waals interactions were shifted between 0.9-1.1 nm. The systems were first energy minimised by steepest descent algorithm to 1000 kJ mol^-1^ nm^-1^ and then simulated for a total of 1 μs. The temperature and pressure were kept constant throughout the simulation at 310 K and 1 bar respectively, with protein, lipids and water/ions coupled individually to a temperature bath by the V-rescale method^54^ and a semi-isotropic Parrinello-Rahman barostat^55^. The final snapshots from the CG simulations were then converted back to an atomistic description using CG2AT^56^.

### Atomistic simulations

The charged N- and C-termini of the converted protein were capped using acetyl and methyl moieties, respectively. All ionisable groups were simulated with default protonation states, unless otherwise mentioned. The virtual site model for hydrogen atoms^57^, adapted for the charmm36 forcefield^58^ was employed, allowing the use of a 4 fs timestep during the simulations. Electrostatics were described using PME, with a cut-off of 1.2 nm and the van der Waals interactions were shifted between 1-1.2 nm. The tip3p water model was used, the water bond angles and distances were constrained by SETTLE^59^. All other bonds were constrained using the LINCS algorithm^60^. The systems were then equilibrated for a further 1 ns using a 4 fs timestep with positional restraints of 1000 kJ mol^-1^ nm^-2^ on the heavy atoms, in a NPT ensemble with temperature V-rescale coupling at 310 K^54^ and semi-isotropic Parrinello-Rahman barostat at 1 bar with protein, lipids and water/ions coupled individually^55^. The Production simulations were performed without position restraints for a total of 200 ns and were run in triplicate.

### Motility assay

*E. coli* RP6894 (Δ*motAB*) was transformed with pT12-derived plasmids encoding C-terminal twin-strep tagged MotAB containing point mutations or appropriate controls. Saturated overnight cultures (2 μL) were injected into soft agar plates (0.3% w/v agar in tryptone broth) containing kanamycin (30 μg/mL) plus rhamnose monohydrate (0.5% w/v) and incubated in a humidified chamber for 23 h at 25 °C.

### Pulldowns

*E. coli* MT56 was transformed with pT12-derived plasmids encoding C-terminal twin-strep tagged MotAB harbouring point mutations or appropriate controls. Small scale (5 mL) cultures were grown at 37 °C for 16 h in TB media containing kanamycin (50 μg/mL) and rhamnose monohydrate (0.1% w/v). OD_600_-normalized cell counts were centrifuged at 4,000*g* for 5 min and pellets frozen at −80 °C. After thawing, cells were lysed by resuspension in 200 mM Tris pH 8.0, 300 mM NaCl, 2 mM EDTA plus 30 μg/mL DNase I and 400 μg/mL lysozyme for 30 min, then solubilized in 1.5 % w/v LMNG for 1 h. Insoluble material was removed by centrifugation at 18,000g for 30 min. LMNG-solubilized lysates were added to 5 μg of TBS-prewashed MagStrep XT magnetic beads (IBA Lifesciences) for 1 h with mild shaking. Beads were isolated and washed twice with TBS plus 0.025% w/v LMNG followed by elution with TBS plus 0.025% w/v LMNG and 50 mM D-biotin. Eluates were diluted in SDS-PAGE sample buffer and run on a 4–20% polyacrylamide gel (NuSep). The gel was stained with InstantBlue (Expedeon) to determine the presence of MotA and MotB.

### Reporting summary

#### Data availability

The data that support the findings of this study are available from the corresponding author upon reasonable request. Cryo-EM volumes and atomic models have been deposited to the EMDB (accession codes EMD-10895, EMD-10899, EMD-10901, EMD-10902, EMD-10897) and PDB (accession codes 6YSF and 6YSL), respectively. Source data for gel shown in Extended Data Fig. 10 are provided with the paper.

## Extended data figures and tables

**Extended Data Fig. 1.**
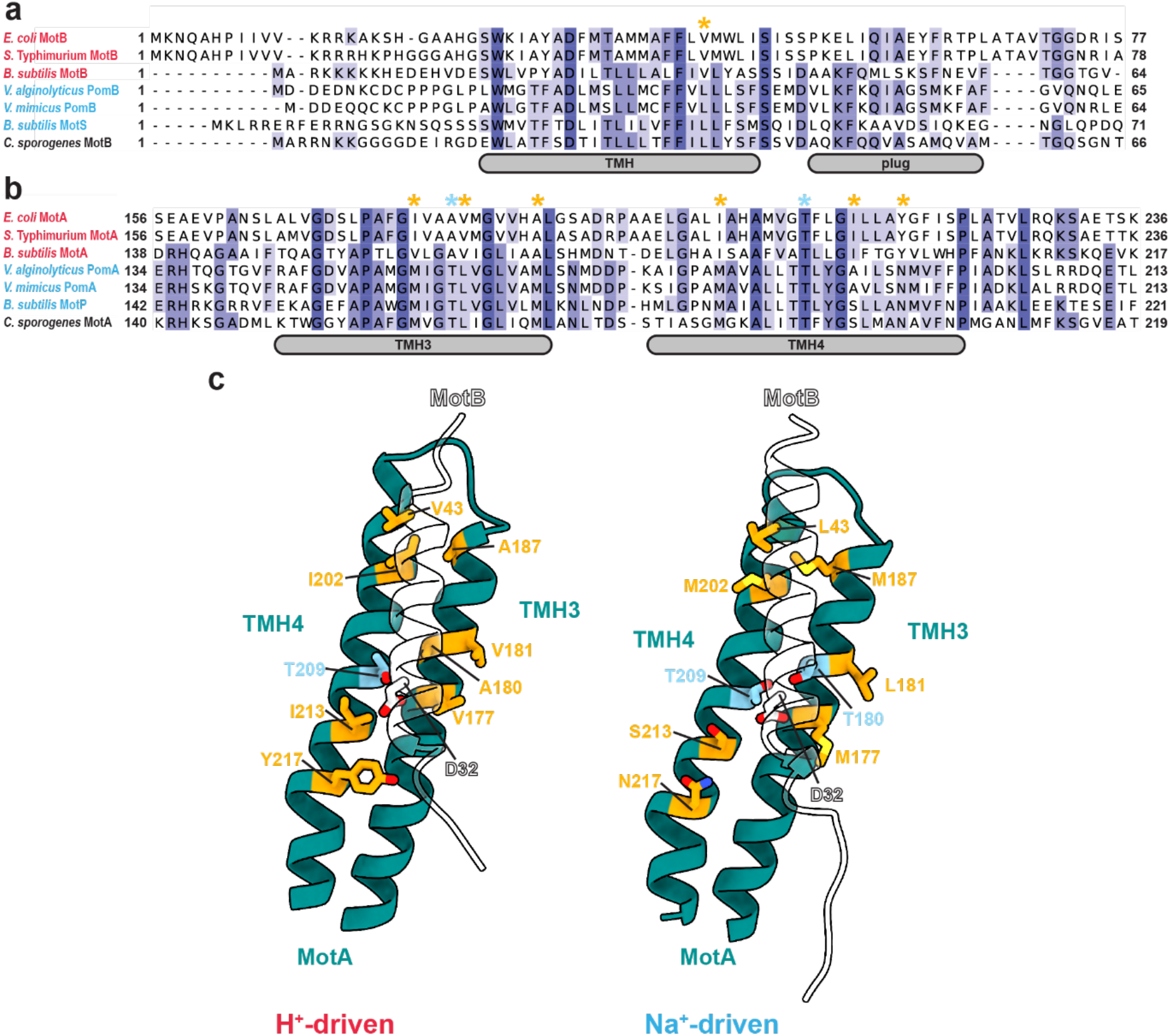
Ion specificity of stator complexes. **a-c,** Sequence alignment of the (a) transmembrane and plug helices of MotB and (b) transmembrane helices (TMHs) 3 and 4 of MotA from *E. coli* and other relevant bacterial species. Species are classified based on ion specificity (red, H^+^-driven; blue, Na^+^-driven). Residues that are conserved just within either the H^+^ or Na^+^-driven classes of stator and map to the MotA-MotB interface (c) are indicated with orange asterisks in the alignment and same residue colouring across H^+^-driven (Left) and Na^+^-driven (Right) models (using *E. coli* MotAB numbering equivalents). The threonine ring is highlighted with light blue asterisks (a,b) or residues (c).

**Extended Data Fig. 2.**
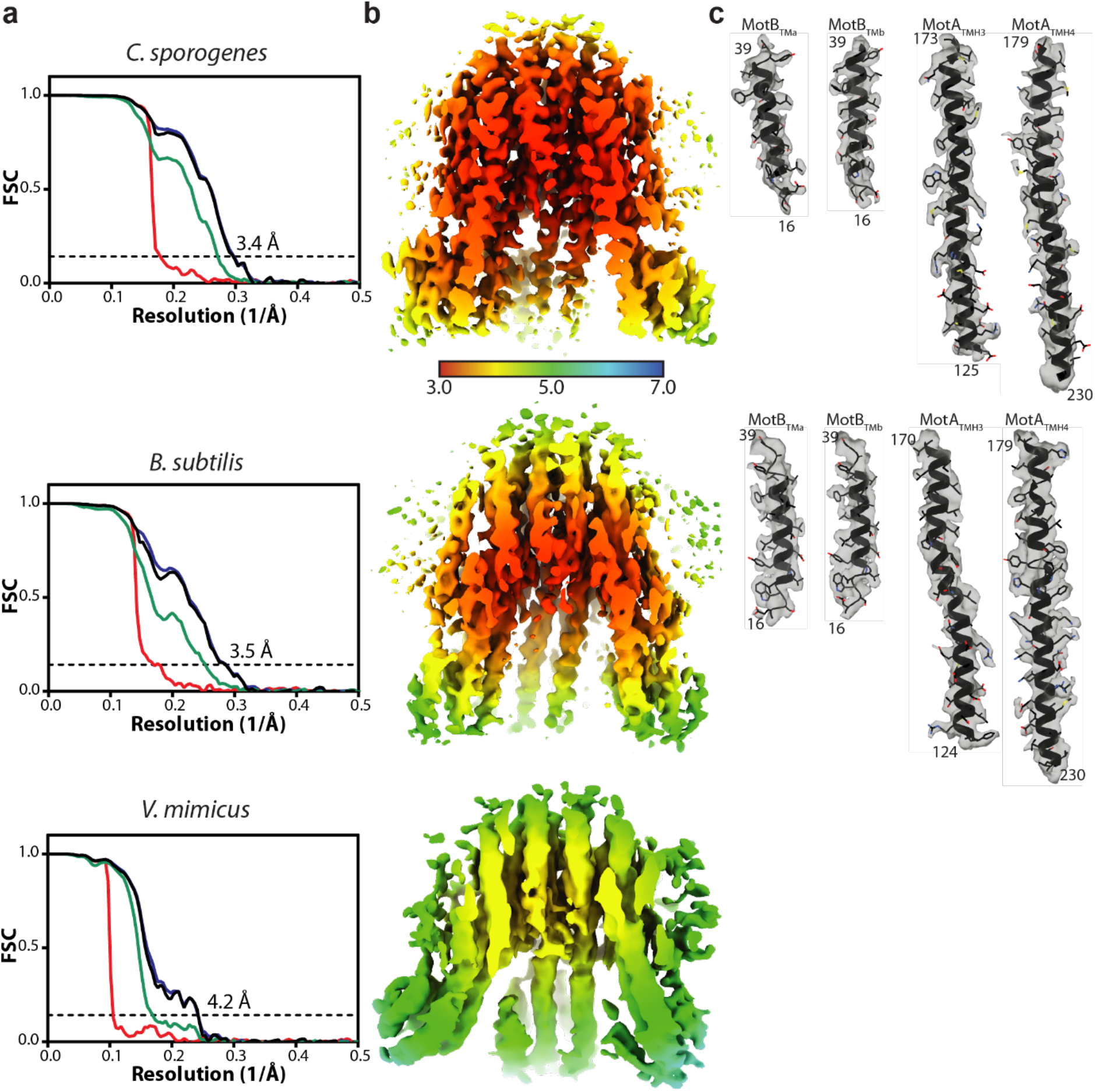
Cryo-EM map quality and resolution estimates of stators. **a**, Gold-standard Fourier shell correlation (FSC) curves of RELION-postprocessed stator volumes. Resolution at the gold-standard cutoff (FSC = 0.143) is indicated. Curves: red, phase-randomized; green, unmasked; blue, masked; black, MTF-corrected. **b**, Local resolution estimates (in Å) of the sharpened volumes. **c**, Representative modelled densities.

**Extended Data Fig. 3.**
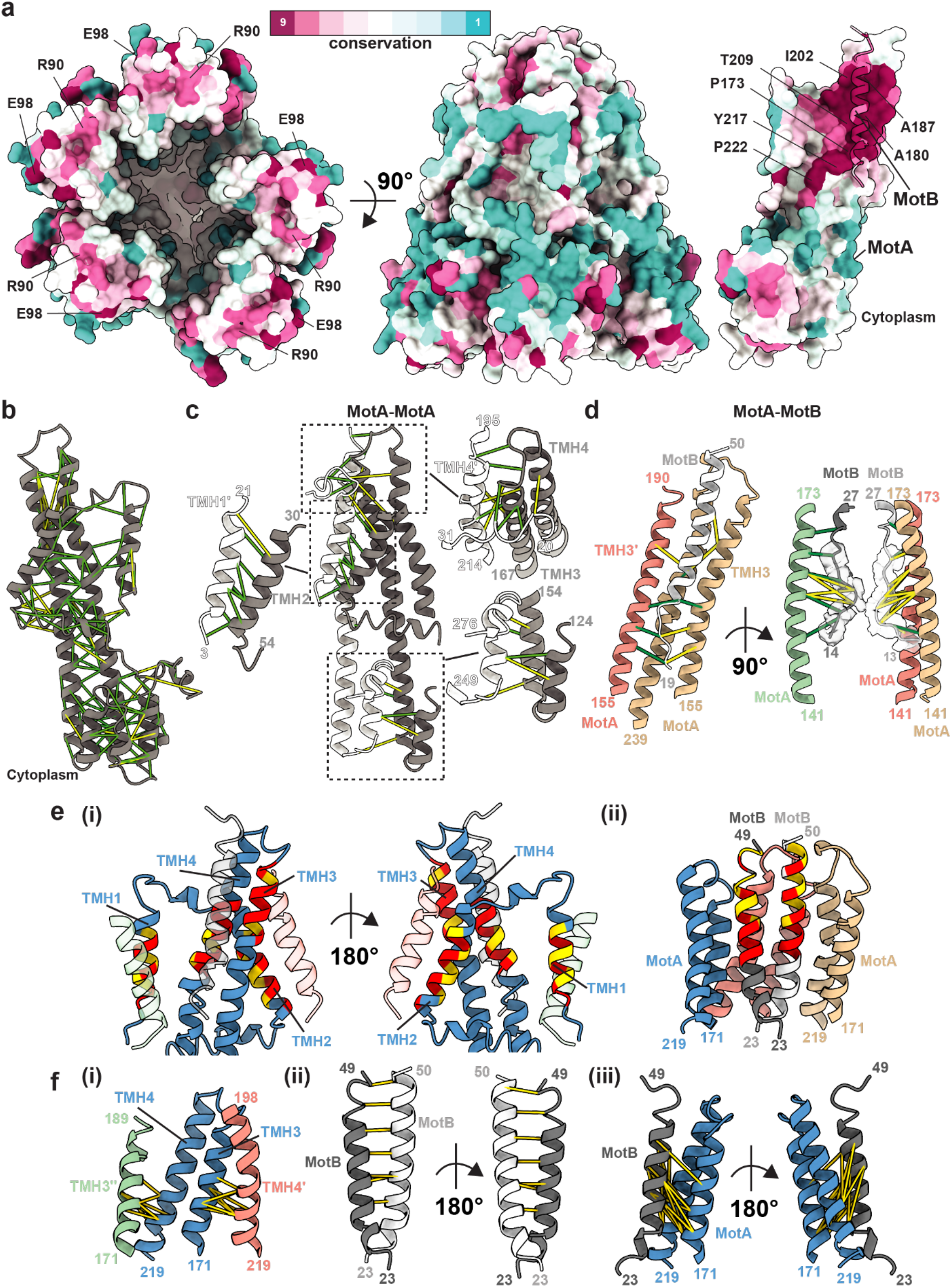
Conservation, covariance, and prior mutagenesis data mapped onto the stator structure. **a**, Surface conservation as determined by Consurf^48^ (maroon high conservation, cyan low conservation). (Left) View from the cytoplasm showing conservation at the cytoplasmic MotA domains, including residues previously identified to be important for torque generation^24^ (R90 and E98). (Centre) Side view showing poor conservation within membrane-interfacing residues of MotA. (Right) Cutaway displaying high level of conservation at the MotA-MotB interface; MotA shown as surface representation, MotB as ribbon. **b-d**, Evolutionary co-variation of residues (b) within MotA, (c) between MotA subunits, with boxes highlighting regions of strong covariance that are illustrated in more detail in the adjacent fragments, or (d) between MotA and MotB with the left hand side showing contacts for modelled MotB regions and the right hand side showing contacts in the unmodelled N-terminal MotB density represented here as a poly-alanine backbone. Predictions were carried out in Gremlin^5^ and contacts with a probability score of > 0.9 are shown. Contacts are coloured by Cα-Cα distance (≤10 Å in green, ≤15 Å in yellow). **e**, Mapping previous tryptophan scanning mutagenesis performed on MotA TMHs^22^ (i) or MotB^23^ (ii) to the stator structure. MotA and MotB are coloured as in Fig. 2,3 with targeted residues coloured according to toleration to mutagenesis; yellow corresponds to tolerated mutants (relative swarm rates > 0.5), red are poorly tolerated mutants (relative swarm rates of ≤ 0.5). In (i), TMH4 (green) and TMH2 (red) of neighbouring MotA subunits and MotB (white) are shown as transparent silhouettes. Poorly tolerated mutants cluster at subunit interfaces. In (ii) only TMH3-TMH4 of three MotA subunits are shown for clarity. **f**, Mapping previously determined cysteine crosslinks between (i) MotA-MotA^20^, (ii) MotB-MotB^21^, or (iii) MotA-MotB^20^ to our structure. For displaying crosslinks, a yield of ≥ 30% disulfide-linked adduct under iodine oxidizing conditions was used as threshold, except for MotA_TMH4_-MotB crosslinks which used a ≥ 10% threshold. All analyses in this figure were performed using an *E. coli* MotAB structure generated by homology threading onto *C. sporogenes* MotAB.

**Extended Data Fig. 4.**
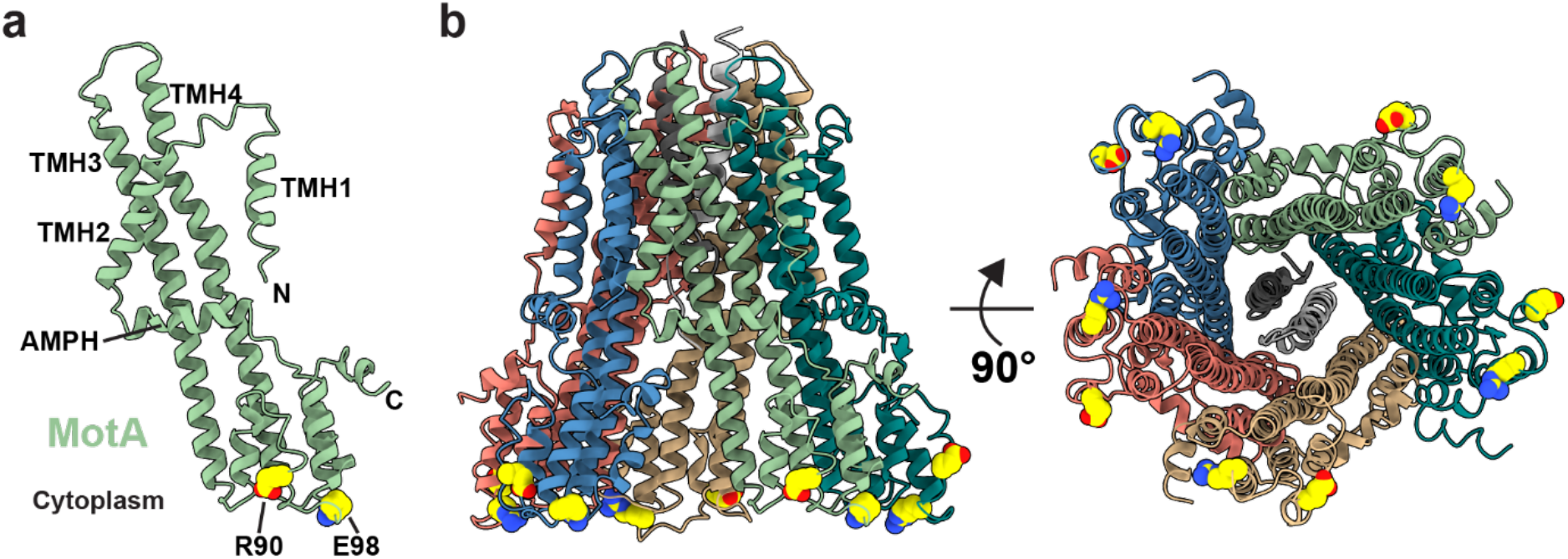
Residues that interact with the flagellar C-ring form a charged ring on the cytoplasmic face of MotA. **a,** An isolated MotA subunit with the essential torque-generating charged residues R90 and E98^24^ displayed in yellow spheres representation. **b,** The full *C. sporogenes* stator complex viewed from the side (Left) or from the cytoplasm (Right), coloured as in Figs. 2-3, and with the torque-generating charged residues represented as in (a) using *E. coli* MotAB numbering scheme.

**Extended Data Fig. 5.**
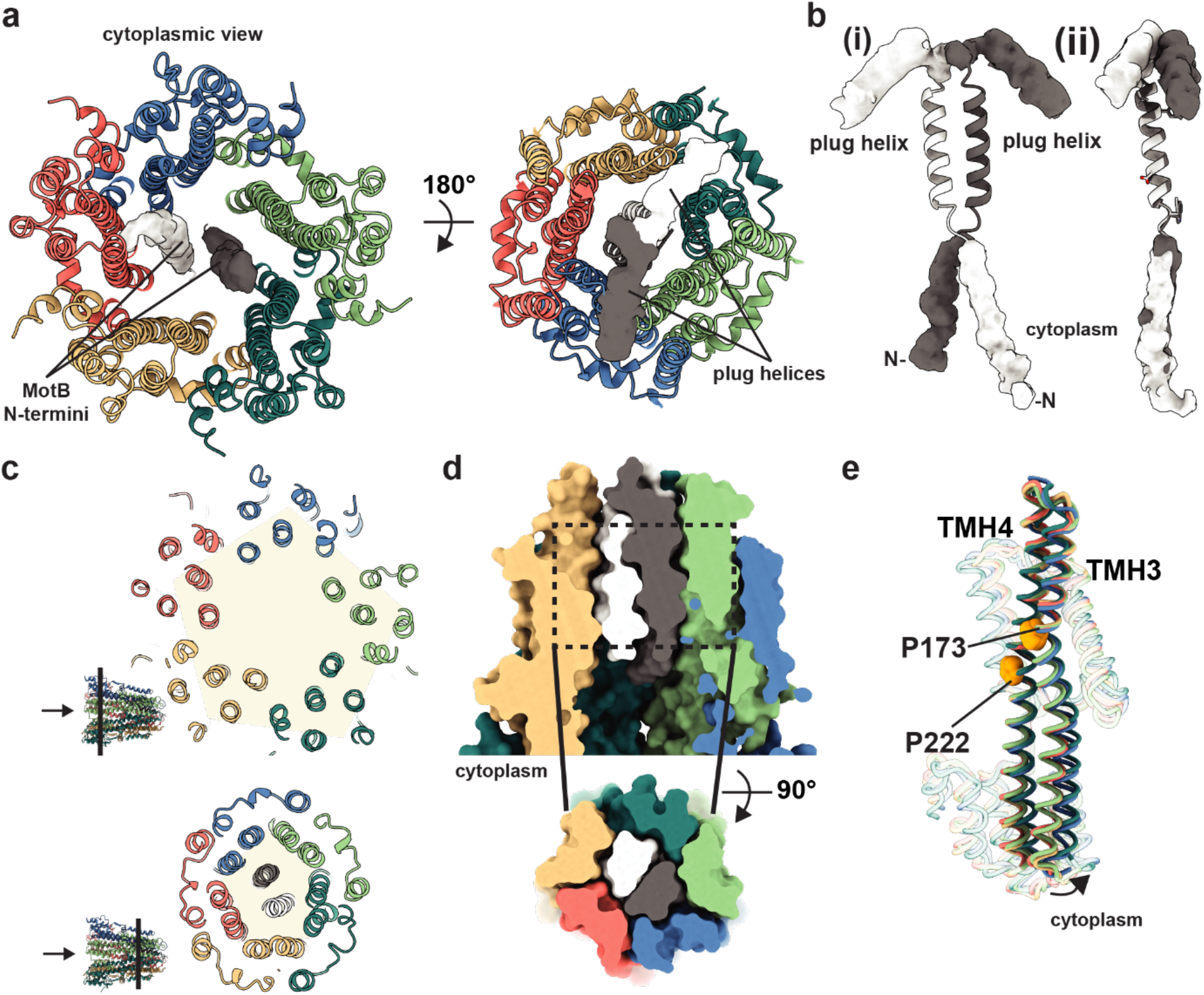
Structural elements of the *B. subtilis* stator. **a**, Structure of the *B. subtilis* stator depicting unmodeled density for the MotB N-terminal extensions (Left) and plug helices (Right). **b**, (i) Isolated *B. subtilis* MotB dimer represented as in (a), and (ii) superposition of the TMHs of the two MotB chains showing the relative rotation of the plug helices. **c**, Slabs at the heights indicated through *B. subtilis* MotAB show distortion from a regular pentagon (arrow indicates the cytoplasmic side of the complex) as viewed from the cytoplasm). **d**, Surface representation of *B. subtilis* MotAB showing tight packing. (Top) Side view with the front of the complex removed. (Bottom) Top-down view of the slab indicated by dashed lines. **e**, Structural alignment of the five *B. subtilis* MotA chains reveal they fall into two conformational classes which differ in the degree of flexing at the highlighted prolines.

**Extended Data Fig. 6.**
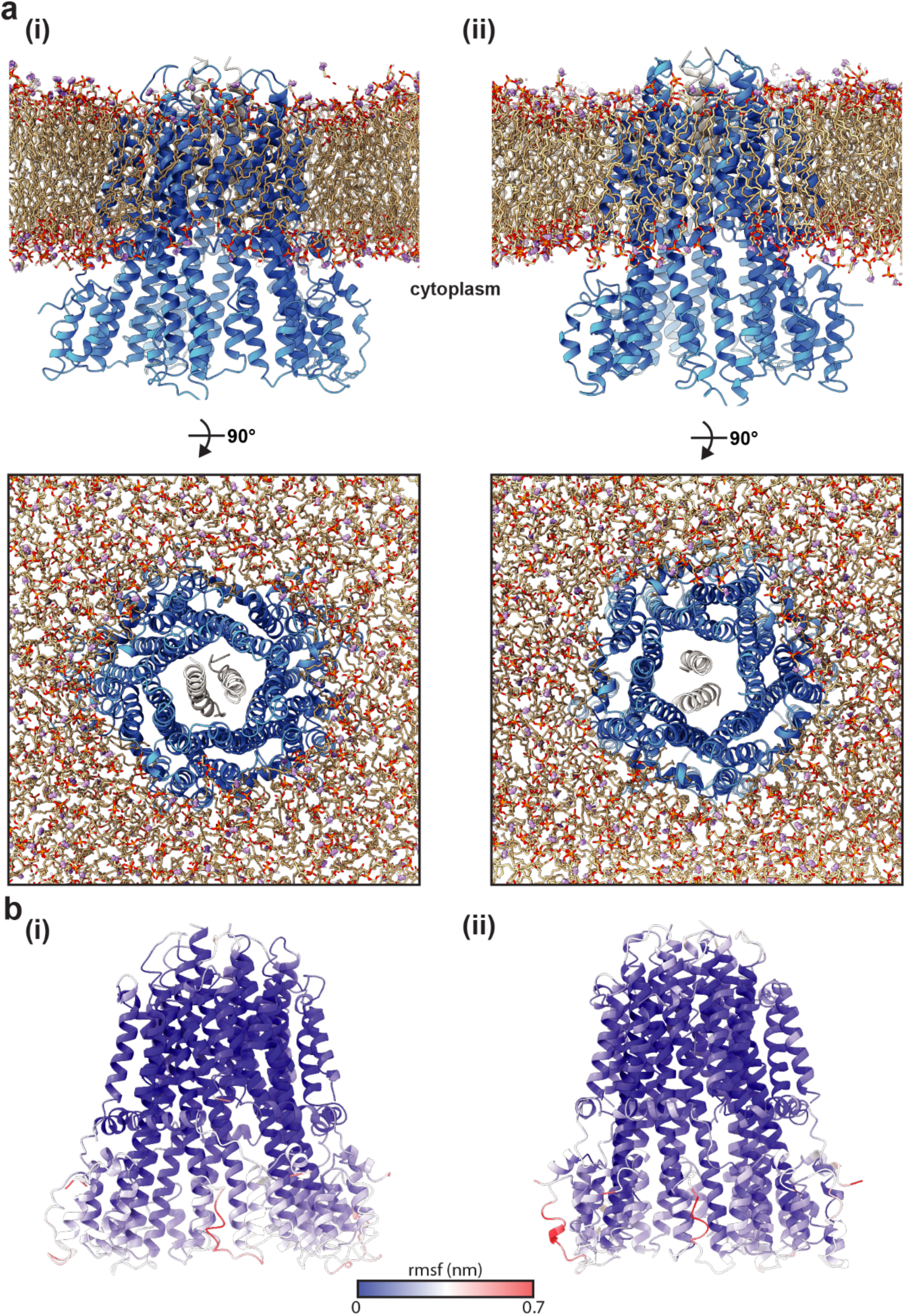
Molecular dynamics simulations of stator structures in lipid bilayers. **a**, Side (*top*) and top-down (*bottom*) views of (i) *C. sporogenes* and (ii) *B. subtilis* MotAB inserted within a lipid bilayer after extended simulations (coarsegrain for 1 μs then atomistic for a further 200 ns). **b**, Cartoon representation of (i) *Clostridium* MotAB and (ii) *Bacillus* MotAB coloured (blue to red) by the average rmsf of the 3 replica simulations performed.

**Extended Data Fig. 7.**
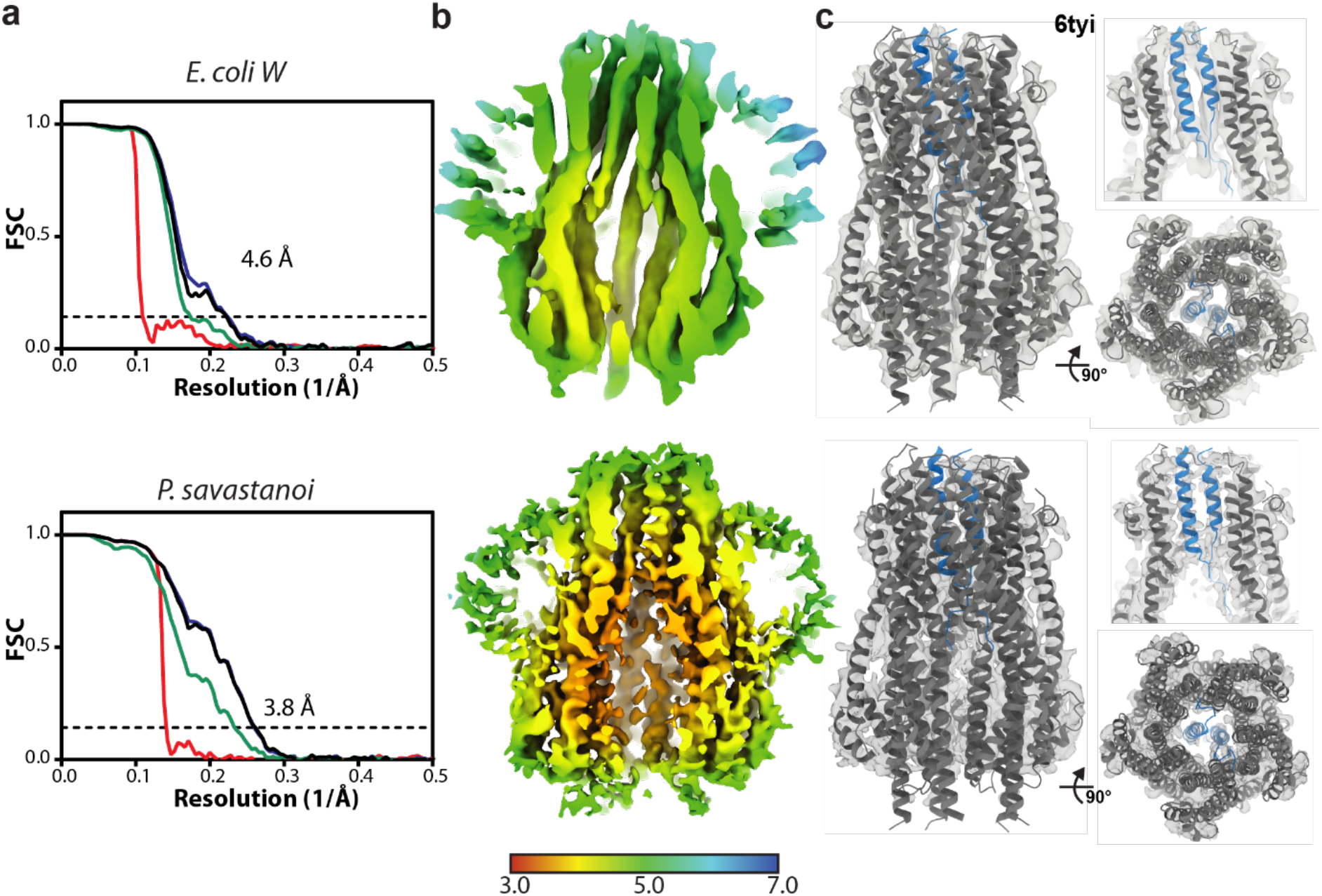
Cryo-EM map quality and resolution estimates of ExbBD complexes. **a**, Gold-standard Fourier shell correlation (FSC) curves of RELION-postprocessed ExbBD maps. Resolution at the gold-standard cutoff (FSC = 0.143) is indicated. Curves: red, phase-randomized; green, unmasked; blue, masked; black, MTF-corrected. **b**, Local resolution estimates (in Å) of the sharpened maps. **c**, Structure of the 5:2 ExbBD complex from *E. coli*^35^ (PDB 6tyi) fit into the *E. coli* W (top) and *P. savastanoi* ExbBD (bottom) maps. ExbB is coloured dark grey and ExbD is coloured blue. Top right panels have three ExbB subunits removed to demonstrate density for the two TMHs of ExbD. Bottom right panels reveal the 5:2 ExbB:ExbD arrangement as viewed from the cytoplasm.

**Extended Data Fig. 8.**
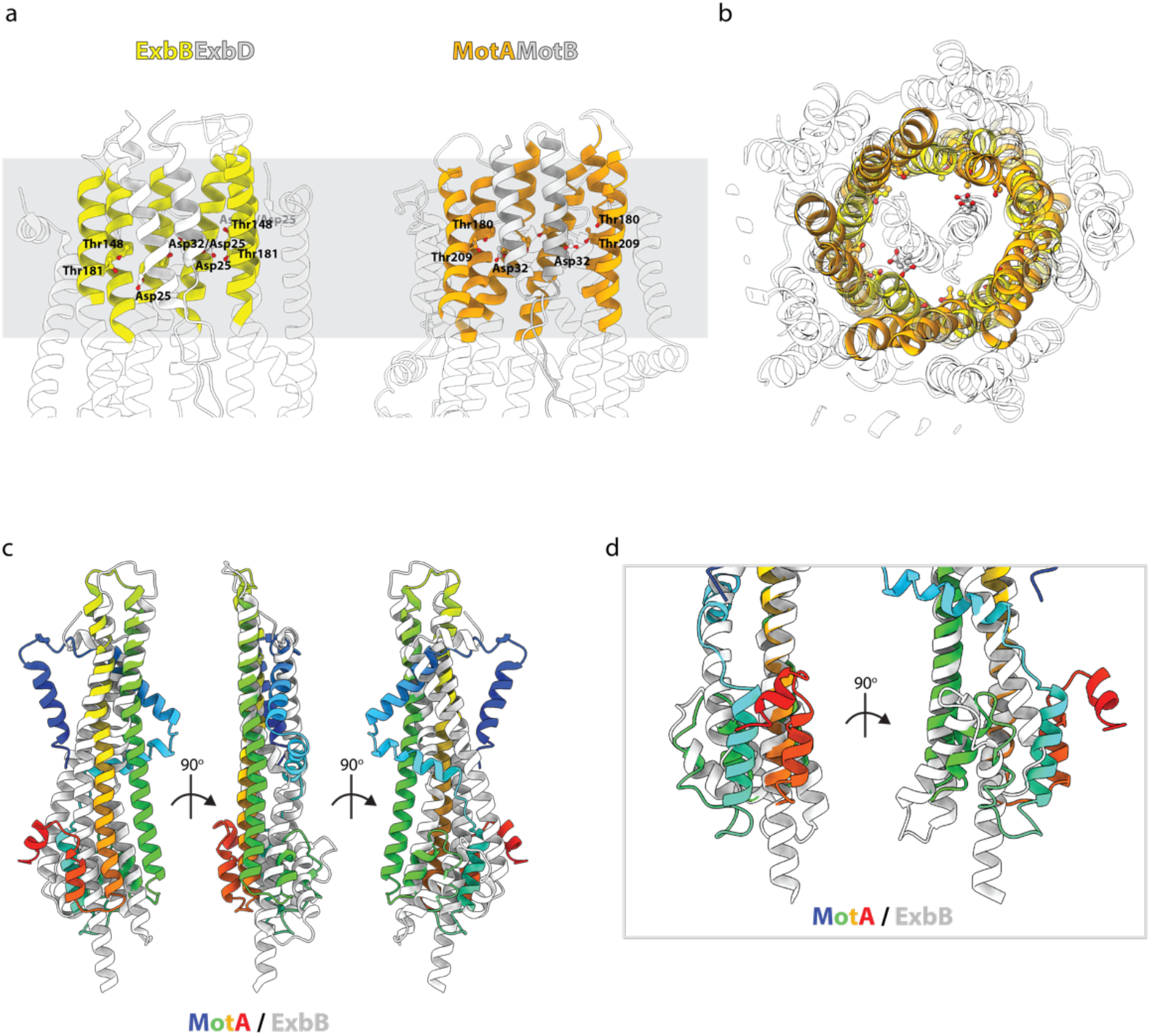
Structural alignment of MotAB and ExbBD. **a,** side views of the cores of the *C. sporogenes* MotAB (orange) and *E. coli* ExbBD^35^ (yellow) complexes with two front MotA/ExbB subunits removed showing the conserved Asp residues lying at the same height with respect to a ring of Thr on the MotA/ExbD components **b,** Overlaying the cores of *C. sporogenes* MotAB (orange) and *E. coli* ExbBD (yellow) by alignment of the MotA/ExbB helices demonstrates alignment of the critical polar residues between the two systems. Regions outside the core shown as transparent grey; MotB/ExbD shown in grey. **c,** Aligning a single subunit of MotA (rainbow colouring) with ExbB (white) via the pore lining helices reveals the different elaborations of this core unit in each protein. **d,** Aligning a single subunit of MotA with ExbB via the cytoplasmic extensions of the two core helices reveals the different folding of the rest of the cytoplasmic regions. Proteins coloured as in (**c**).

**Extended Data Fig. 9.**
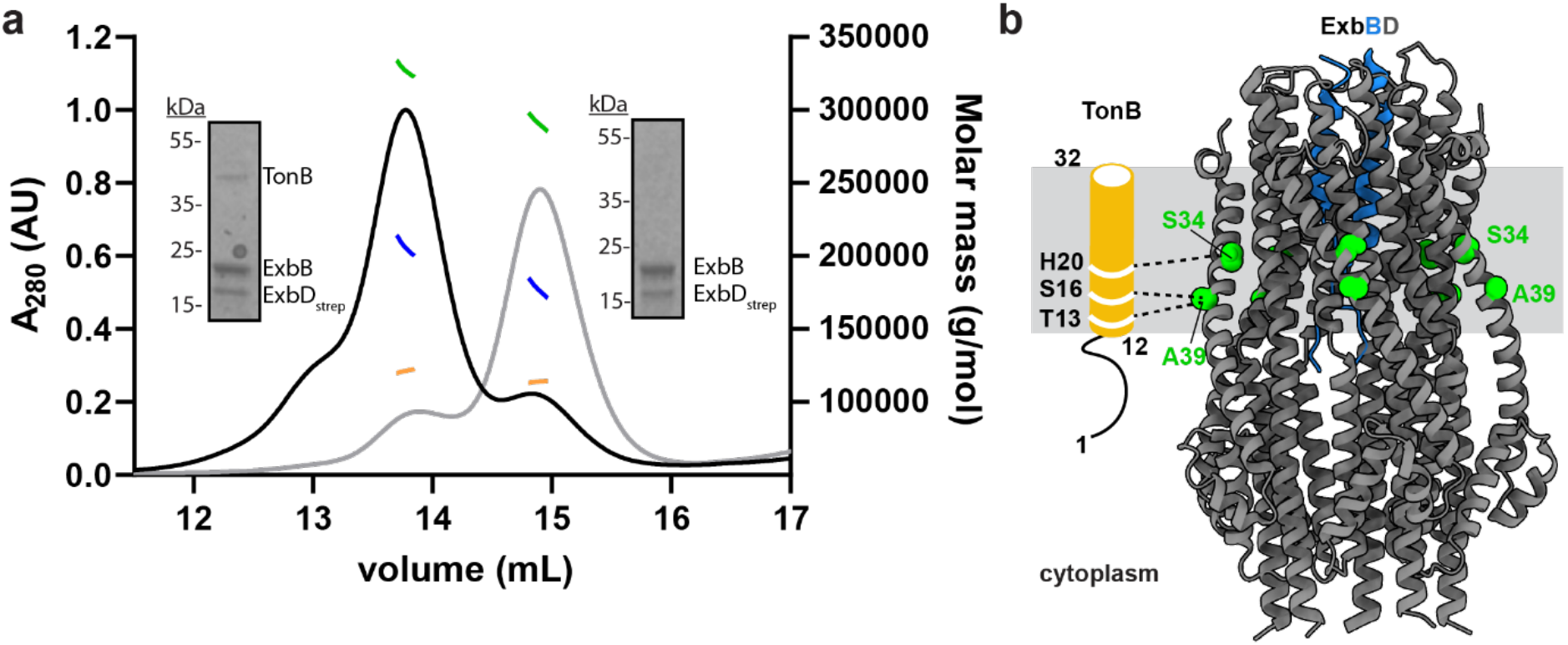
TonB recruitment to the ExbBD complex. **a,** SEC-MALS profile of purified *Pseudomonas* TonB-ExbBD (black absorption curve) and ExbBD (grey absorption curve) with SDS-PAGE analysis of each sample inlayed. Total protein-detergent complex molar mass (green) and deconvoluted protein (blue) and detergent (orange) molar masses are shown. A ~30 kDa difference in molar mass is observed between the complexes consistent with the the TonB-ExbBD complex containing one TonB subunit. **b,** Evolutionary co-variation of residues between TonB and ExbB displayed on the *E. coli* ExbBD structure^35^ (PDB 6tyi). For clarity, a topological model of the TMH of TonB is shown in orange with TonB-ExbB contacts indicated as dashed lines. Covarying residues decorating the periphery of ExbB are displayed in green. ExbB (grey) and ExbD (blue) are displayed as ribbon cartoons. Contacts shown were generated by Gremlin^5^ and have a probability score of > 0.9.

## Supplementary Data

**Supplementary Table 1.** Bacterial strains and plasmids used in this study

**Supplementary Table 2.** Oligonucleotides used in this study.

**Supplementary table 3.** Species conversion of residues specified in text

**Supplementary Table 4.** List of evolutionary contacts predicted by Gremlin

## References

1 Berg, H. C. The rotary motor of bacterial flagella. Annu Rev Biochem 72, 19–54, doi:10.1146/annurev.biochem.72.121801.161737 (2003).

2 Leeuwenhoek, A. Observation, communicated to the publisher by Mr Anthony van Leewenhoeck, in a Dutch letter of the 9 Octob. 1676 here English’d: concerning little animals by him observed in rain-well-sea and snow water; as also in water wherein pepper had lain infused. Phil. Trans. 12, 821–831, doi:doi:10.1098/rstl.1677.0003 (1677).

3 Nakamura, S. & Minamino, T. Flagella-Driven Motility of Bacteria. Biomolecules 9, doi:10.3390/biom9070279 (2019).

4 Johnson, S. et al. Symmetry mismatch in the MS-ring of the bacterial flagellar rotor explains the structural coordination of secretion and rotation. Nat Microbiol, doi:10.1038/s41564-020-0703-3 (2020).

5 Ovchinnikov, S. et al. Large-scale determination of previously unsolved protein structures using evolutionary information. Elife 4, e09248, doi:10.7554/eLife.09248 (2015).

6 Kojima, S. Dynamism and regulation of the stator, the energy conversion complex of the bacterial flagellar motor. Curr Opin Microbiol 28, 66–71, doi:10.1016/j.mib.2015.07.015 (2015).

7 Blair, D. F. & Berg, H. C. Restoration of torque in defective flagellar motors. Science 242, 1678–1681, doi:10.1126/science.2849208 (1988).

8 Blair, D. F. & Berg, H. C. The MotA protein of E. coli is a proton-conducting component of the flagellar motor. Cell 60, 439–449, doi:10.1016/0092-8674(90)90595-6 (1990).

9 Larsen, S. H., Adler, J., Gargus, J. J. & Hogg, R. W. Chemomechanical coupling without ATP: the source of energy for motility and chemotaxis in bacteria. Proc Natl Acad Sci US A 71, 1239–1243, doi:10.1073/pnas.71.4.1239 (1974).

10 Asai, Y., Yakushi, T., Kawagishi, I. & Homma, M. Ion-coupling determinants of Na+-driven and H+-driven flagellar motors. J Mol Biol 327, 453–463, doi:10.1016/s0022-2836(03)00096-2 (2003).

11 Kojima, S. & Blair, D. F. Solubilization and purification of the MotA/MotB complex of Escherichia coli. Biochemistry 43, 26–34, doi:10.1021/bi035405l (2004).

12 Nirody, J. A., Sun, Y.-R. & Lo, C.-J. The biophysicist’s guide to the bacterial flagellar motor. Advances in Physics: X 2, 324–343, doi:10.1080/23746149.2017.1289120 (2017).

13 Kojima, S. et al. The Helix Rearrangement in the Periplasmic Domain of the Flagellar Stator B Subunit Activates Peptidoglycan Binding and Ion Influx. Structure 26, 590–598 e595, doi:10.1016/j.str.2018.02.016 (2018).

14 Zhu, S. et al. Conformational change in the periplamic region of the flagellar stator coupled with the assembly around the rotor. Proc Natl Acad Sci U S A 111, 13523–13528, doi:10.1073/pnas.1324201111 (2014).

15 Kim, E. A., Price-Carter, M., Carlquist, W. C. & Blair, D. F. Membrane segment organization in the stator complex of the flagellar motor: implications for proton flow and proton-induced conformational change. Biochemistry 47, 11332–11339, doi:10.1021/bi801347a (2008).

16 Kojima, S. & Blair, D. F. Conformational change in the stator of the bacterial flagellar motor. Biochemistry 40, 13041–13050, doi:10.1021/bi011263o (2001).

17 Mandadapu, K. K., Nirody, J. A., Berry, R. M. & Oster, G. Mechanics of torque generation in the bacterial flagellar motor. Proc Natl Acad Sci U S A 112, E4381–4389, doi:10.1073/pnas.1501734112 (2015).

18 Boschert, R., Adler, F. R. & Blair, D. F. Loose coupling in the bacterial flagellar motor. Proc Natl Acad Sci U S A 112, 4755–4760, doi:10.1073/pnas.1419955112 (2015).

19 Nishihara, Y. & Kitao, A. Gate-controlled proton diffusion and protonation-induced ratchet motion in the stator of the bacterial flagellar motor. Proc Natl Acad Sci U S A 112, 7737–7742, doi:10.1073/pnas.1502991112 (2015).

20 Braun, T. F., Al-Mawsawi, L. Q., Kojima, S. & Blair, D. F. Arrangement of core membrane segments in the MotA/MotB proton-channel complex of Escherichia coli. Biochemistry 43, 35–45, doi:10.1021/bi035406d (2004).

21 Braun, T. F. & Blair, D. F. Targeted disulfide cross-linking of the MotB protein of Escherichia coli: evidence for two H(+) channels in the stator Complex. Biochemistry 40, 13051–13059, doi:10.1021/bi011264g (2001).

22 Sharp, L. L., Zhou, J. & Blair, D. F. Features of MotA proton channel structure revealed by tryptophan-scanning mutagenesis. Proc Natl Acad Sci U S A 92, 7946–7950, doi:10.1073/pnas.92.17.7946 (1995).

23 Sharp, L. L., Zhou, J. & Blair, D. F. Tryptophan-scanning mutagenesis of MotB, an integral membrane protein essential for flagellar rotation in Escherichia coli. Biochemistry 34, 9166–9171, doi:10.1021/bi00028a028 (1995).

24 Yakushi, T., Yang, J., Fukuoka, H., Homma, M. & Blair, D. F. Roles of charged residues of rotor and stator in flagellar rotation: comparative study using H+-driven and Na+-driven motors in Escherichia coli. J Bacteriol 188, 1466–1472, doi:10.1128/JB.188.4.1466-1472.2006 (2006).

25 Hosking, E. R. & Manson, M. D. Clusters of charged residues at the C terminus of MotA and N terminus of MotB are important for function of the Escherichia coli flagellar motor. J Bacteriol 190, 5517–5521, doi:10.1128/JB.00407-08 (2008).

26 Hosking, E. R., Vogt, C., Bakker, E. P. & Manson, M. D. The Escherichia coli MotAB proton channel unplugged. J Mol Biol 364, 921–937, doi:10.1016/j.jmb.2006.09.035 (2006).

27 Braun, T. F. et al. Function of proline residues of MotA in torque generation by the flagellar motor of Escherichia coli. J Bacteriol 181, 3542–3551 (1999).

28 Minamino, T., Kinoshita, M. & Namba, K. Directional Switching Mechanism of the Bacterial Flagellar Motor. Comput Struct Biotechnol J 17, 1075–1081, doi:10.1016/j.csbj.2019.07.020 (2019).

29 Lam, K. H. et al. Multiple conformations of the FliG C-terminal domain provide insight into flagellar motor switching. Structure 20, 315–325, doi:10.1016/j.str.2011.11.020 (2012).

30 Khan, S., Dapice, M. & Humayun, I. Energy transduction in the bacterial flagellar motor. Effects of load and pH. Biophys J 57, 779–796, doi:10.1016/S0006-3495(90)82598-4 (1990).

31 Manson, M. D., Tedesco, P. M. & Berg, H. C. Energetics of flagellar rotation in bacteria. J Mol Biol 138, 541–561, doi:10.1016/s0022-2836(80)80017-9 (1980).

32 Marmon, L. Elucidating the origin of the ExbBD components of the TonB system through Bayesian inference and maximum-likelihood phylogenies. Mol Phylogenet Evol 69, 674–686, doi:10.1016/j.ympev.2013.07.010 (2013).

33 Celia, H. et al. Structural insight into the role of the Ton complex in energy transduction. Nature 538, 60–65, doi:10.1038/nature19757 (2016).

34 Maki-Yonekura, S. et al. Hexameric and pentameric complexes of the ExbBD energizer in the Ton system. Elife 7, doi:10.7554/eLife.35419 (2018).

35 Celia, H. et al. Cryo-EM structure of the bacterial Ton motor subcomplex ExbB-ExbD provides information on structure and stoichiometry. Commun Biol 2, 358, doi:10.1038/s42003-019-0604-2 (2019).

36 Swayne, C. & Postle, K. Taking the Escherichia coli TonB transmembrane domain “offline”? Nonprotonatable Asn substitutes fully for TonB His20. J Bacteriol 193, 3693–3701, doi:10.1128/JB.05219-11 (2011).

37 Ollis, A. A., Kumar, A. & Postle, K. The ExbD periplasmic domain contains distinct functional regions for two stages in TonB energization. J Bacteriol 194, 3069–3077, doi:10.1128/JB.00015-12 (2012).

38 Pawelek, P. D. et al. Structure of TonB in complex with FhuA, E. coli outer membrane receptor. Science 312, 1399–1402, doi:10.1126/science.1128057 (2006).

39 Cascales, E., Lloubes, R. & Sturgis, J. N. The TolQ-TolR proteins energize TolA and share homologies with the flagellar motor proteins MotA-MotB. Mol Microbiol 42, 795–807, doi:10.1046/j.1365-2958.2001.02673.x (2001).

40 McBride, M. J. & Zhu, Y. Gliding motility and Por secretion system genes are widespread among members of the phylum bacteroidetes. J Bacteriol 195, 270–278, doi:10.1128/JB.01962-12 (2013).

41 Kuhlen, L. et al. Structure of the core of the type III secretion system export apparatus. Nat Struct Mol Biol 25, 583–590, doi:10.1038/s41594-018-0086-9 (2018).

42 Zivanov, J., Nakane, T. & Scheres, S. H. W. A Bayesian approach to beam-induced motion correction in cryo-EM single-particle analysis. IUCrJ 6, 5–17, doi:10.1107/S205225251801463X (2019).

43 Rohou, A. & Grigorieff, N. CTFFIND4: Fast and accurate defocus estimation from electron micrographs. J Struct Biol 192, 216–221, doi:10.1016/j.jsb.2015.08.008 (2015).

44 Reboul, C. F., Eager, M., Elmlund, D. & Elmlund, H. Single-particle cryo-EM-Improved ab initio 3D reconstruction with SIMPLE/PRIME. Protein Sci 27, 51–61, doi:10.1002/pro.3266 (2018).

45 Brown, A. et al. Tools for macromolecular model building and refinement into electron cryo-microscopy reconstructions. Acta Crystallogr D Biol Crystallogr 71, 136–153, doi:10.1107/S1399004714021683 (2015).

46 Afonine, P. V. et al. Real-space refinement in PHENIX for cryo-EM and crystallography. Acta Crystallogr D Struct Biol 74, 531–544, doi:10.1107/S2059798318006551 (2018).

47 Williams, C. J. et al. MolProbity: More and better reference data for improved all-atom structure validation. Protein Sci 27, 293–315, doi:10.1002/pro.3330 (2018).

48 Ashkenazy, H. et al. ConSurf 2016: an improved methodology to estimate and visualize evolutionary conservation in macromolecules. Nucleic Acids Res 44, W344–350, doi:10.1093/nar/gkw408 (2016).

49 Kelley, L. A., Mezulis, S., Yates, C. M., Wass, M. N. & Sternberg, M. J. The Phyre2 web portal for protein modeling, prediction and analysis. Nat Protoc 10, 845–858, doi:10.1038/nprot.2015.053 (2015).

50 Goddard, T. D. et al. UCSF ChimeraX: Meeting modern challenges in visualization and analysis. Protein Sci 27, 14–25, doi:10.1002/pro.3235 (2018).

51 Abraham, M. J. et al. GROMACS: High performance molecular simulations through multi-level parallelism from laptops to supercomputers. SoftwareX 1-2, 19–25, doi:10.1016/j.softx.2015.06.001 (2015).

52 de Jong, D. H. et al. Improved Parameters for the Martini Coarse-Grained Protein Force Field. Journal of Chemical Theory and Computation 9, 687–697, doi:10.1021/ct300646g (2012).

53 Wassenaar, T. A., Ingólfsson, H. I., Böckmann, R. A., Tieleman, D. P. & Marrink, S. J. Computational Lipidomics with insane: A Versatile Tool for Generating Custom Membranes for Molecular Simulations. Journal of Chemical Theory and Computation 11, 2144–2155, doi:10.1021/acs.jctc.5b00209 (2015).

54 Bussi, G., Donadio, D. & Parrinello, M. Canonical sampling through velocity rescaling. The Journal of Chemical Physics 126, doi:10.1063/1.2408420 (2007).

55 Parrinello, M. & Rahman, A. Polymorphic transitions in single crystals: A new molecular dynamics method. Journal of Applied Physics 52, 7182–7190, doi:10.1063/1.328693 (1981).

56 Stansfeld, P. J. & Sansom, M. S. P. From Coarse Grained to Atomistic: A Serial Multiscale Approach to Membrane Protein Simulations. Journal of Chemical Theory and Computation 7, 1157–1166, doi:10.1021/ct100569y (2011).

57 Feenstra, K. A., Hess, B. & Berendsen, H. J. C. Improving efficiency of large time-scale molecular dynamics simulations of hydrogen-rich systems. Journal of Computational Chemistry 20, 786–798, doi:10.1002/(sici)1096-987x(199906)20:8<786::Aid-jcc5>3.0.Co;2-b (1999).

58 Olesen, K., Awasthi, N., Bruhn, D. S., Pezeshkian, W. & Khandelia, H. Faster Simulations with a 5 fs Time Step for Lipids in the CHARMM Force Field. Journal of Chemical Theory and Computation 14, 3342–3350, doi:10.1021/acs.jctc.8b00267 (2018).

59 Miyamoto, S. & Kollman, P. A. Settle: An analytical version of the SHAKE and RATTLE algorithm for rigid water models. Journal of Computational Chemistry 13, 952–962, doi:10.1002/jcc.540130805 (1992).

60 Hess, B., Bekker, H., Berendsen, H. J. C. & Fraaije, J. G. E. M. LINCS: A linear constraint solver for molecular simulations. Journal of Computational Chemistry 18, 1463–1472, doi:10.1002/(sici)1096-987x(199709)18:12<1463::Aid-jcc4>3.0.Co;2-h (1997).

